# TMEM41B is a pan-flavivirus host factor

**DOI:** 10.1101/2020.10.09.334128

**Authors:** H.-Heinrich Hoffmann, William M Schneider, Kathryn Rozen-Gagnon, Linde A Miles, Felix Schuster, Brandon Razooky, Eliana Jacobson, Xianfang Wu, Soon Yi, Charles M Rudin, Margaret R MacDonald, Laura K McMullan, John T Poirier, Charles M Rice

**Author notes:** These authors contributed equally to this work (H.-Heinrich Hoffmann, William M Schneider). These authors contributed equally to this work (Linde A Miles, Felix Schuster, Brandon Razooky, Eliana Jacobson).

## Abstract

Flaviviruses pose a constant threat to human health. These RNA viruses are transmitted by the bite of infected mosquitoes and ticks and regularly cause outbreaks. To identify host factors required for flavivirus infection we performed full-genome loss of function CRISPR-Cas9 screens. Based on these results we focused our efforts on characterizing the roles that TMEM41B and VMP1 play in the virus replication cycle. Our mechanistic studies on TMEM41B revealed that all members of the *Flaviviridae* family that we tested require TMEM41B. We tested 12 additional virus families and found that SARS-CoV-2 of the *Coronaviridae* also required TMEM41B for infection. Remarkably, single nucleotide polymorphisms (SNPs) present at nearly twenty percent in East Asian populations reduce flavivirus infection. Based on our mechanistic studies we hypothesize that TMEM41B is recruited to flavivirus RNA replication complexes to facilitate membrane curvature, which creates a protected environment for viral genome replication.

**HIGHLIGHTS:** TMEM41B and VMP1 are required for both autophagy and flavivirus infection, however, autophagy is not required for flavivirus infection.

TMEM41B associates with viral proteins and likely facilitates membrane remodeling to establish viral RNA replication complexes.

TMEM41B single nucleotide polymorphisms (SNPs) present at nearly twenty percent in East Asian populations reduce flavivirus infection.

TMEM41B-deficient cells display an exaggerated innate immune response upon high multiplicity flavivirus infection.

## INTRODUCTION

Virus outbreaks are a regular, yet unpredictable feature of an interconnected world. Viruses of the *Flaviviridae* family are known to cause outbreaks worldwide and are transmitted primarily by the bite of infected mosquitoes and ticks.

Flaviviruses, members of the *Flaviviridae* family, are positive sense single-stranded RNA viruses that have caused several notable outbreaks in recent history. For example, West Nile virus (WNV) emerged in New York City in 1999, spread across the continent, and is now endemic in the United States (Kramer et al., 2019; Roehrig et al., 2002). Also noteworthy are the recurring yellow fever virus (YFV) outbreaks that occur in sub-Saharan Africa and South America despite the availability of a highly effective vaccine (Ahmed and Memish, 2017). Most recently, the 2016 Zika virus (ZIKV) epidemic swept through South and Central America wreaking havoc on a generation of unborn children by causing microcephaly *in utero* (Hills et al., 2017; Lee and Ng, 2018). In addition to these outbreaks, *Aedes albopictus*, the natural vector for ZIKV, dengue virus (DENV) and chikungunya virus (CHIKV), has been found consistently in recent years in Central Europe (Muller et al., 2020), and the prevalence of tick-borne diseases is at a record high. These alarming trends correlate with warming temperatures due to climate change, which expand the range of mosquitoes and ticks (Brady and Hay, 2020; Brugueras et al., 2020; Dobler, 2010; McPherson et al., 2017; Medlock et al., 2013). These recent flavivirus outbreaks underscore the devastating impacts viral outbreaks have on human health and the constant need for preparedness.

For many viruses with the potential to cause outbreaks there are no specific treatments. In instances where antiviral therapies do exist, they are often virus specific. One strategy to prepare for and respond to viral outbreaks is to develop drugs that target host factors viruses require to complete their replication cycles. CRISPR-Cas9 gene disruption has been used to identify host factors required for virus infection and nominate candidate drug targets. We, like others, set out to identify factors required for Zika virus as well as other flaviviruses. Here we present our screen results and detailed studies focused on two autophagy related genes, transmembrane protein 41B (*TMEM41B)* and vacuole membrane protein 1 (*VMP1*), that were enriched in our screens.

Autophagy, or “self-eating”, is a conserved multistep process in which cytoplasmic and intracellular components are engulfed in double membrane structures called autophagosomes. Following a series of maturation steps, autophagosomes ultimately fuse with lysosomes where their contents are degraded and recycled (Dikic and Elazar, 2018). We found that the full autophagy pathway is not required for flavivirus infection; however, TMEM41B and VMP1 are essential. In comparison to VMP1, little is known about the cellular role of TMEM41B, and no data have been reported about the role of TMEM41B in flavivirus infection. Here we show that TMEM41B is a pan-flavivirus host factor that is required for all flaviviruses we tested which included both mosquito-borne and tick-borne flaviviruses. Beyond the classical flaviviruses, we also found that TMEM41B is required by additional members within the *Flaviviridae* including hepatitis C virus (HCV) from the hepacivirus genus and bovine viral diarrhea virus (BVDV) from the pestivirus genus. We further tested a broad panel of viruses representing 12 virus families including negative- and positive-sense RNA viruses, retroviruses and DNA viruses, and found that of these, the only other virus that required TMEM41B was SARS-CoV-2 of the *Coronaviridae*. Indeed, we recently identified *TMEM41B* as a top scoring hit in multiple coronavirus genome-wide CRISPR-Cas9 loss of function screens, including screens with SARS-CoV-2 (see accompanying manuscript (Schneider et al., 2020)). It is possible that coronaviruses and flaviviruses require TMEM41B for a similar step in their replication cycles. The mechanistic studies we present here demonstrates that TMEM41B relocalizes upon flavivirus infection and associates with viral proteins, nonstructural protein 4A (NS4A) and NS4B, which are known to form membrane-bound viral RNA replication complexes (Neufeldt et al., 2018; Paul and Bartenschlager, 2015). These and other data we present suggest that TMEM41B is likely hijacked during the flavivirus replication cycle to facilitate the membrane remodeling necessary for replication of viral RNA.

## RESULTS

### Genome-wide CRISPR-Cas9 screens for Zika and yellow fever viruses identify TMEM41B and VMP1 as required host factors

To identify host factors required for ZIKV and YFV, we performed pooled genome-wide CRISPR knockout (KO) screens in B3GALT6-deficient human haploid (HAP1) cells. We used B3GALT6 KO cells to potentially decrease cellular heparan sulfate (HS) protein glycosylation which has been shown to nonspecifically facilitate virus binding and subsequent entry (Cagno et al., 2019). We hypothesized that this alteration in surface protein glycosylation would decrease the enrichment and overabundance of hits related to HS protein glycosylation and nonspecific virus binding. Nevertheless, consistent with a wide range of virus-selected genome-wide loss of function screens (Hoffmann et al., 2017; Jae et al., 2014; Realegeno et al., 2017; Riblett et al., 2016), we identified host factors involved in heparan sulfate biosynthesis.

Like other flavivirus host factor screens, we identified genes that are involved in oligosaccharide transfer, and genes involved in protein translocation and folding into the endoplasmic reticulum (ER) (**Figure 1A**) (Marceau et al., 2016; Savidis et al., 2016; Zhang et al., 2016). Outside of these categories, *TMEM41B* was among the top scoring genes. Little is known about the cellular function of TMEM41B; however, three independent genome-wide CRISPR-Cas9 loss of function screens identified *TMEM41B* as a gene that plays an important role in autophagy (Moretti et al., 2018; Morita et al., 2019; Shoemaker et al., 2019). These groups went on to show that TMEM41B is functionally similar to VMP1, which is known to have a role in lipid mobilization and autophagy (Morishita et al., 2019; Ropolo et al., 2007; Zhao et al., 2017).

**Figure 1.**
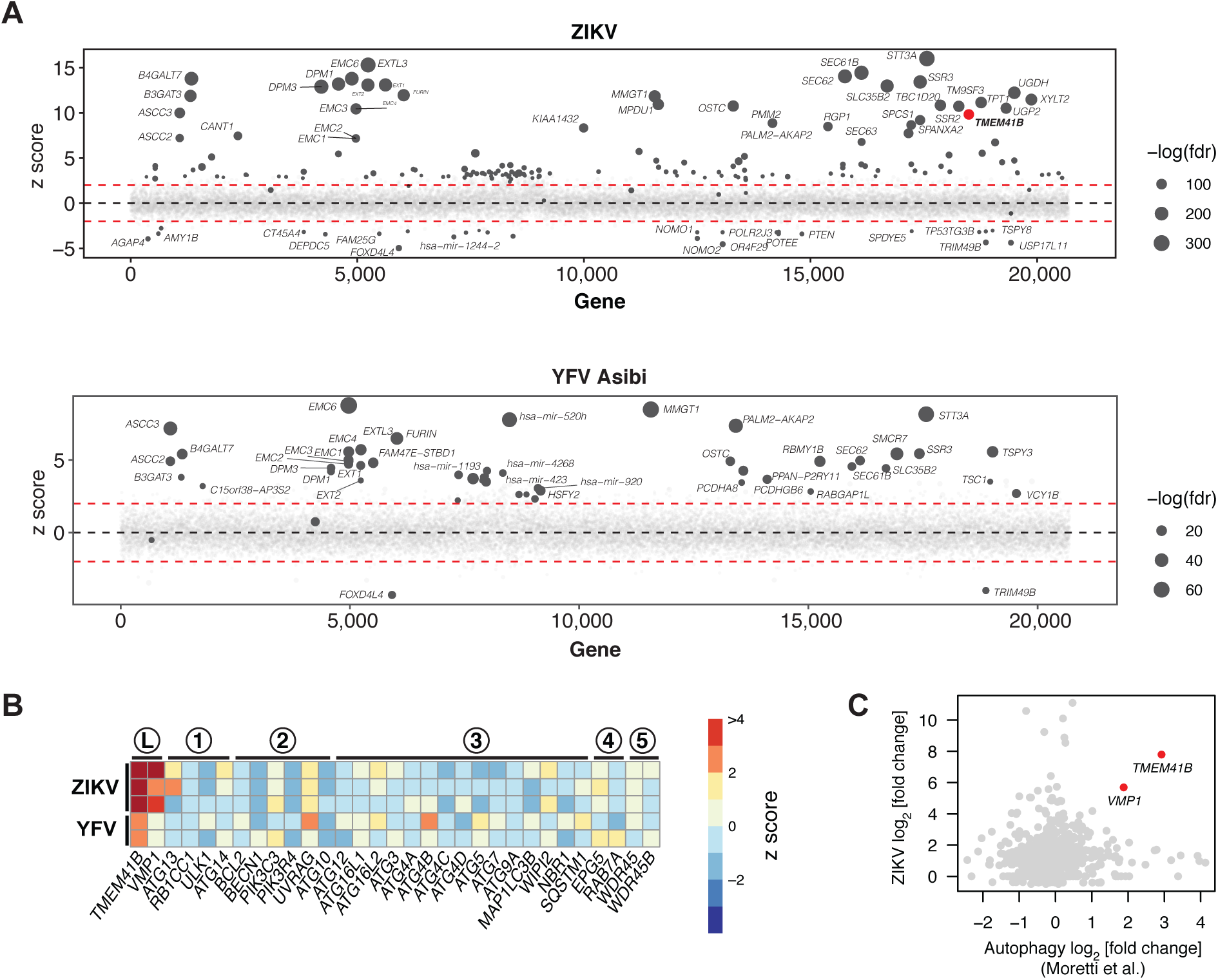
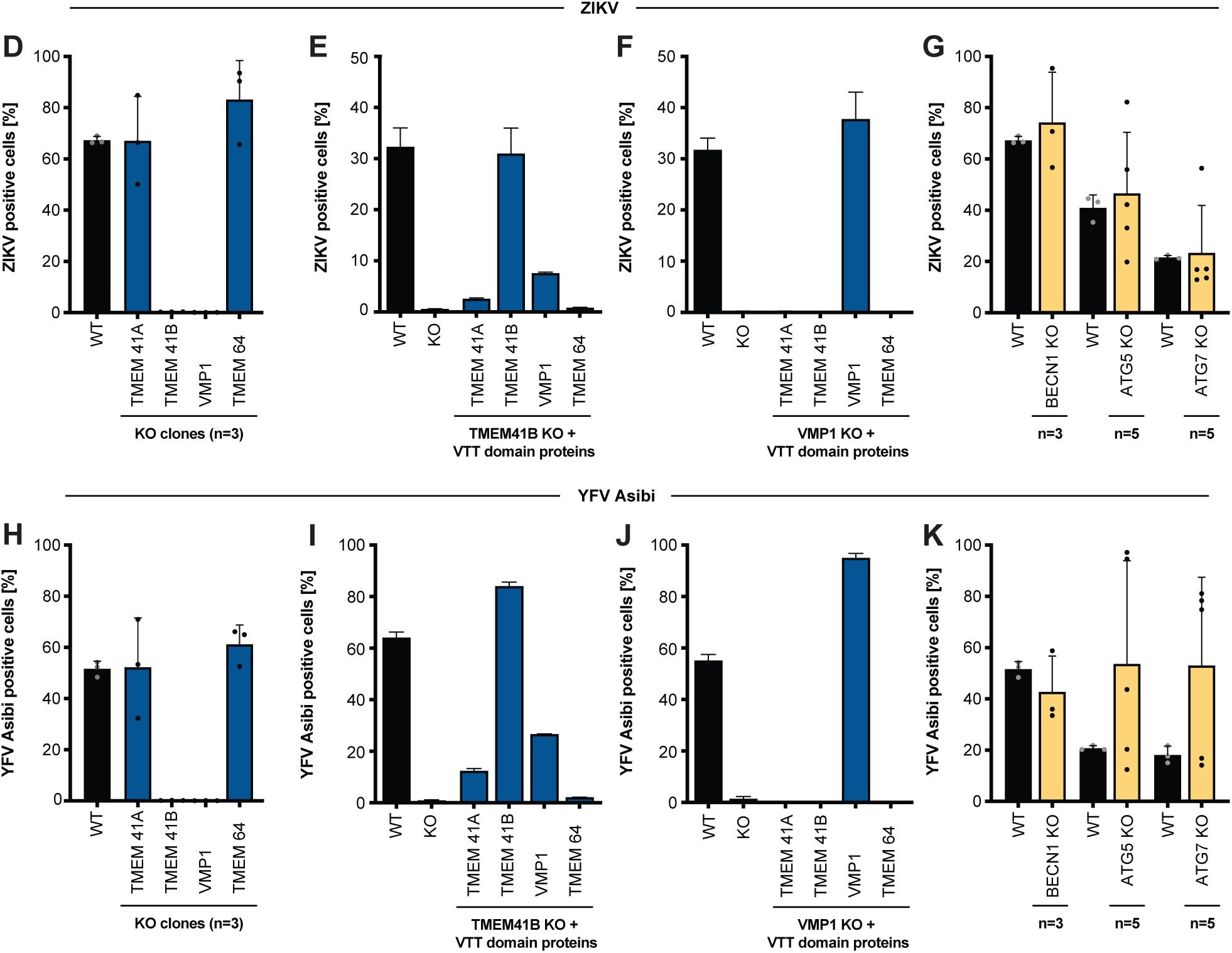
Genome-wide CRISPR-Cas9 screens for Zika and yellow fever viruses identify TMEM41B and VMP1 as required host factors. **(A)** Bubble plot of data from ZIKV screen (top) and YFV screen (bottom). Red lines denote z = ± 2. **(B)** Heatmap of z scores for genes in the autophagy pathway ordered sequentially by functional role: L, lipid mobilization; 1, initiation; 2, nucleation; 3, elongation; 4, sequestration; 5, tethering/fusion. **(C)** Scatter plot of gene-wise log2 fold change (LFC) from this study (ZIKV) versus Moretti et al. autophagy screen. **(D)** HAP1 wildtype (WT) and (n=3) individual knockout (KO) clones for VTT domain-containing proteins infected with ZIKV (MOI = 0.5 PFU/cell) for 48 h. **(E)** WT and TMEM41B KO HAP1 cells overexpressing individual VTT domain proteins infected with ZIKV (MOI = 0.25 PFU/cell) for 48 h. **(F)** WT and VMP1 KO HAP1 cells overexpressing individual VTT domain proteins infected with ZIKV ((MOI = 0.25 PFU/cell) for 48 h. **(G)** HAP1 WT and (n=3-5) individual KO clones for autophagy genes infected with ZIKV (MOI = 0.5 PFU/cell) for 48 h. **(H-K)** Same as panels D-G but infected with YFV Asibi (MOI = 0.1 PFU/cell) for 72 h. Cells were analyzed by flow cytometry and plotted as a percentage of viral antigen positive cells. Dots in panels D, G, H, and K represent the average of n=3 replicates from individual single cell clones. Error bars in panels E, F, I, and J depict standard deviation (SD) of n = three replicates. See also Figure S1 B-I.

There are numerous, sometimes conflicting reports, which indicate that autophagy-related genes can promote or restrict *Flaviviridae* infection. This literature has been recently reviewed by Po-Yuan Ke (Ke, 2018). Our identification of TMEM41B prompted us to interrogate our screen data further for genes involved in autophagy. Of a list of genes with an established role in autophagy, only *TMEM41B* and *VMP1*, which act at the early stage of autophagy, scored positively (**Figure 1B**). This list, however, is not comprehensive, and therefore we next compared our ZIKV screen results to those from a genome-wide CRISPR-Cas9 screen that was designed to identify novel genes required for autophagy (Moretti et al., 2018). As shown in **Figure 1C**, these two screens showed no overlap with the exception of two genes: *TMEM41B* and *VMP1*.

### The VTT domain proteins TMEM41B and VMP1 are required for flavivirus infection

TMEM41B and VMP1 are multipass transmembrane proteins that share a stretch of amino acids recently named the VTT domain based on homology among <u>V</u>MP1, <u>T</u>MEM41A/B, and <u>T</u>MEM64 (Morita et al., 2019). Given this homology and the fact that both TMEM41B and VMP1 were top scoring hits in our flavivirus screens, we sought to test all four members of the VTT domain family for their requirement in flavivirus infection. Further, in addition to ZIKV and YFV (strain: Asibi) used in our screens, we expanded our collection of flaviviruses for these experiments to include three additional flaviviruses: the YFV 17D (vaccine strain), DENV-2, and WNV.

To test which of the four VTT domain-containing proteins are required for flavivirus infection, we used CRISPR-Cas9 gene disruption to generate clonal HAP1 cell lines individually lacking VMP1, TMEM41A, TMEM41B, and TMEM64 (**Figure S1A**). We then infected three to five independent knockout (KO) clones for each of these candidate host factors with each of the five flaviviruses listed above or with human parainfluenza virus 3 (hPIV-3) as a non-flavivirus control. Consistent with our screening results, we found that of these four VTT domain-containing proteins, only TMEM41B and VMP1 are required for flavivirus infection as cells deficient for these two proteins do not support virus replication (**Figure 1D, H** and **S1B, D, F**). While TMEM41B was not required for hPIV-3 infection (**Figure S1H**), decreased growth of hPIV-3 was observed in the VMP1 KO clones. VMP1 KO clones appear less viable and grow slower compared to WT cells, which likely accounts for the slight decrease in replication seen for the control virus, hPIV-3 (**Figure S1H**).

We next expressed each of the four VTT domain-containing proteins individually in TMEM41B KO and VMP1 KO cells to test their ability to restore virus infection. As expected, TMEM41B overexpression completely restored infection in TMEM41B KO cells. Interestingly, we found that VMP1 and TMEM41A overexpression could also partially compensate for the lack of TMEM41B during flavivirus infection; however TMEM41A was less efficient at restoring infection compared to VMP1. In contrast, only VMP1 was able to rescue infection in VMP1 KO cells (**Figure 1E, F, I, J**). Furthermore, VMP1 expression restored the aforementioned observed decrease in cell viability in VMP1 KO cells and also rescued hPIV-3 replication (**Figure S1J, K**). The capacity of VMP1 to rescue a TMEM41B defect but not vice versa is consistent with what has been shown for TMEM41B and VMP1 in the context of autophagy (Morita et al., 2019).

### Classical autophagy proteins BECN1, ATG5, and ATG7 are not required for flavivirus infection in HAP1 cells

Since both TMEM41B and VMP1 have been shown to be involved in autophagy, we hypothesized that other canonical autophagy related genes could be essential for flavivirus infection (Moretti et al., 2018; Morita et al., 2019; Ropolo et al., 2007; Shoemaker et al., 2019). To test this hypothesis, we generated clones lacking BECN1, ATG5, and ATG7, which are known to play a critical role in autophagy (Kang et al., 2011; Mizushima et al., 1998; Tanida et al., 1999). We confirmed KO in these clones by genome sequencing as well as by western blot (for ATG5 and ATG7) (**Figure S1L**). While ATG5 and ATG7 KO clones already lack lipidated LC3 II at baseline levels (indicative of defective autophagy (Tanida et al., 2008; Tanida and Waguri, 2010)), we sought to test ATG7 KO clones in addition for their ability to respond to classical autophagy stimuli (e.g., starvation and treatment with Torin 1 in the presence of chloroquine). As expected, ATG7-deficient cells are unable to mount an autophagy response. This is shown by western blot in **Figure S1M**, where autophagy results in LC3 I lipidation and conversion to LC3 II in WT cells but not in ATG7 KO HAP1 clones (Tanida et al., 2008; Tanida and Waguri, 2010). We then tested these autophagy-deficient cells for their ability to support flavivirus infection and found that cells lacking BECN1, ATG5, or ATG7 retained their ability to support flavivirus infection (**Figure 1G, K** and **S1C, E, G, I**). These data suggest that autophagy alone is not necessarily required for flavivirus infection. This does not, however, rule out the possibility that non-canonical autophagy mechanisms are required for flavivirus infection nor does it suggest TMEM41B and VMP1 are the only autophagy factors required for flavivirus infection. In fact, two other non-classical autophagy genes were highly enriched in our YFV screen (TBC1D5 and TBC1D20) and in our ZIKV screen (TBC1D20) underscoring the complex interplay between flavivirus replication and the autophagy pathway (Popovic et al., 2012; Popovic and Dikic, 2014; Sidjanin et al., 2016; Sklan et al., 2007) (**Figure 1A, B)**.

### TMEM41B is a pan-flavivirus host factor

We have shown that mosquito-borne flaviviruses require TMEM41B for infection. To gain additional insight into the requirement of this host factor for viral infection, we extended our studies to include tick-borne flaviviruses, viruses from additional genera within the *Flaviviridae* family, and a diverse panel of unrelated viruses.

The tick-borne flaviviruses we tested include Powassan virus (POWV), a BSL3 pathogen currently expanding in North America in *Ixodes* ticks (Dennis et al., 1998; Ebel, 2010; Eisen et al., 2016), and five BSL4 pathogens: two strains of tick-borne encephalitis virus (TBEV) representing the European and Far Eastern clade, and three hemorrhagic fever viruses, Omsk hemorrhagic fever virus (OHFV), Kyasanur forest disease virus (KFDV), and Alkhurma hemorrhagic fever virus (AHFV). In addition, we generated TMEM41B KO clones in hepatocellular carcinoma cells (Huh-7.5) and bovine MBDK cells to test additional members in the *Flaviviridae*: HCV in the hepacivirus genus and BVDV in pestivirus genus. As shown in **Figure 2A-D**, all of these viruses require TMEM41B for infection. This demonstrates that TMEM41B is a pan-flavivirus host factor.

**Figure 2.**
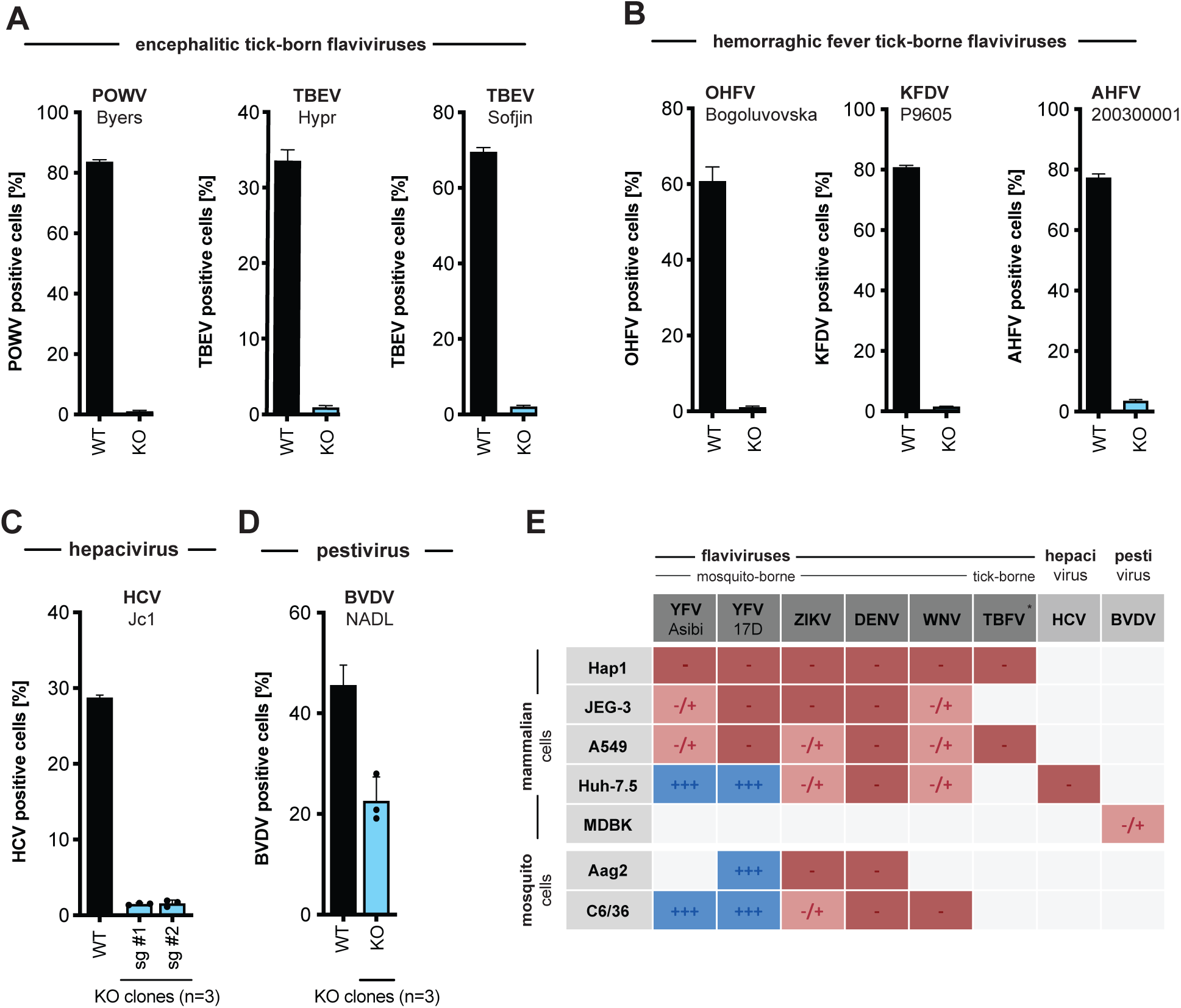
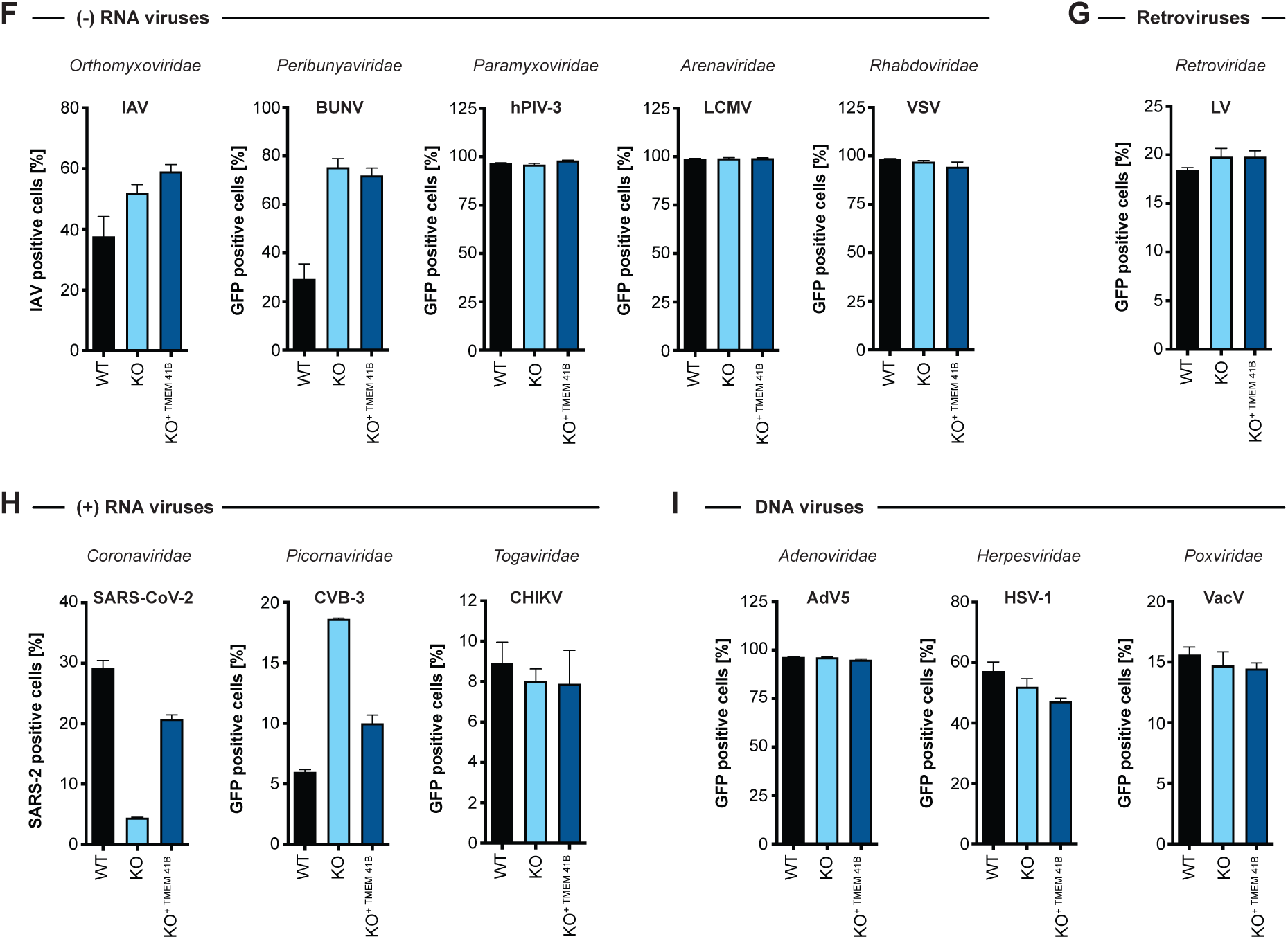
TMEM41B is a pan-flavivirus host factor. **(A)** WT and TMEM41B KO HAP1 cells infected for 48 h with encephalitic tick-borne flaviviruses POWV at MOI = 0.02 PFU/cell, and for 24h with TBEV (strain: Hypr) at MOI = 0.02 PFU/cell, TBEV (strain: Sofjin) at MOI = 0.02 PFU/cell. **(B)** WT and TMEM41B KO HAP1 cells infected for 24 h with hemorrhagic fever tick-borne flaviviruses OHFV at MOI = 0.02 PFU/cell, KFDV at MOI = 0.02 PFU/cell, AHFV at MOI = 0.02 PFU/cell. **(C)** WT and TMEM41B KO Huh-7.5 clones infected with HCV-RFP for 72 h at MOI = 0.05 PFU/cell. **(D)** TMEM41B KO MDBK clones infected with BVDV for 48 h at MOI = 0.01 PFU/cell. Dots represent the average of n=3 replicates from individual single cell clones. **(A-D)** Virus strain is indicated in parenthesis. Error bars in panels A and B show SD for n=6 replicates. Error bars in panels C and D show SD for n=3 replicates (WT) and individual KO clones. **(E)** Table summarizing results shown in Figure S2 (panels A, B, D, F and H-K) from infection experiments with multiple viruses in various mammalian and mosquito cell lines. TBFV = tick-born flaviviruses of panel A and B, +++ = infection comparable to WT cells; +/-= reduced but detectable infection; - = negligible infection. **(F)** WT, TMEM41B KO and reconstituted HAP1 cells infected with (-) sense RNA viruses. Influenza A virus (IAV), MOI = 0.1 PFU/cell for 24 h; bunyamwera virus (BUNV-GFP), MOI = 0.025 IU/cell for 48 h; human parainfluenza virus 3 (h-PIV3-GFP), MOI = 0.02 IU/cell for 48 h; lymphocytic choriomeningitis virus (LCMV-GFP), MOI = 0.01 IU/cell for 48 h; vesicular stomatitis virus (VSV-GFP), MOI = 0.01 PFU/cell for 24 h. **(G)** WT, TMEM41B KO and reconstituted HAP1 cells infected with VSV-G-pseudotyped HIV-1 lentivirus (LV-GFP), MOI = (1:25 dilution of virus stock) for 48 h. **(H)** WT, TMEM41B KO and reconstituted HAP1 cells infected with (+) sense RNA viruses. Cells infected with SARS-CoV-2 were prior reconstituted to stably express ACE2 and TMPRSS2, MOI = 1 PFU/cell for 24 h; Coxsackie virus B (CVB-3-GFP), MOI = (1:100 dilution of virus stock) for 8 h; chikungunya virus (CHIKV-GFP), MOI = 0.025 PFU/cell for 24 h. **(I)** WT, TMEM41B KO and reconstituted HAP1 cells infected with DNA viruses. Adenovirus 5 (AdV5-GFP), MOI = 200 p/cell for 24 h; herpes simplex virus 1 (HSV-1-GFP), MOI = 2.5 PFU/cell for 8 h; vaccinia virus (VacV-GFP), MOI = 0.005 PFU/cell for 48 h. Cells were analyzed by flow cytometry and plotted as a percentage of viral antigen positive cells or GFP/RFP positive cells for reporter viruses expressing a fluorescent protein. Error bars depict SD for three replicates or the indicated number of clones.

Flaviviruses replicate in a variety of tissue types. Therefore, we generated TMEM41B KO and reconstituted cell clones in additional mammalian cell lines representing different tissue types including lung carcinoma (A549) and placenta carcinoma (JEG3) epithelial cells. In addition, we knocked out and reconstituted the mosquito TMEM41B homologue in C6/36 cells (*Aedes albopictus*) and Aag2 cells (*Aedes aegypti*) to determine if TMEM41B was necessary for flavivirus infection in the mosquito vector. We found that TMEM41B was required in nearly all virus-cell combinations (**Figure 2E** and **S2A-L**). However, we observed several virus-cell combinations where TMEM41B appears to be less critical. For example, whereas TMEM41B is critical for YFV infection in HAP1, JEG3, and A549 cells, it is not required in Huh-7.5 cells. Notably we even observed differences in the requirement for TMEM41B among two strains of the same virus: the wildtype Asibi strain of YFV and the 17D vaccine strain that differ by only 31 amino acids (**Figure 1H**, **S1B** and **S2A, D, F**) (dos Santos et al., 1995). Given that VMP1 can compensate for TMEM41B deficiency, we hypothesized that differences in TMEM41B and VMP1 expression among cell types could potentially explain this variability. We therefore performed western blots on HAP1, JEG3, A549, and Huh-7.5 cell lysates to compare VMP1 and TMEM41B abundance. We found that full-length TMEM41B is highly expressed in HAP1 and JEG3 cells, but present at much lower abundance in A549 and Huh-7.5 cells. Further, the TMEM41B antibody detects several abundant lower molecular weight bands in A549 cells that may or may not be TMEM41B isoforms. VMP1, in contrast, is expressed at much lower levels in HAP1 cells compared to JEG3, A549, and Huh-7.5 cells (**Figure S2M**). While speculative, we posit that the differential requirement for TMEM41B that we observe in some virus-cell combinations is due to differences in TMEM41B and VMP1 abundance among cell types and the extent to which VMP1 can compensate for TMEM41B deficiency.

Given that TMEM41B is broadly required among members of the *Flaviviridae*, we posited that other more distantly related viruses or unrelated viruses may share a similar requirement for TMEM41B. Using the TMEM41B KO and reconstituted HAP1 cells, we queried the TMEM41B requirement for a diverse panel of viruses including negative- and positive-sense RNA viruses, retroviruses and DNA viruses. Interestingly, we found that Coxsackievirus 3B (CVB3) infection is *enhanced* in the absence of TMEM41B, whereas infection with SARS-CoV-2, a member of the *Coronaviridae*, is impaired similar to the viruses in the *Flaviviridae,* suggesting that it also requires TMEM41B for infection. Aside from these two viruses, none of the other viruses tested were affected by the lack of TMEM41B (**Figure 2F-I**). Our observation that SARS-CoV-2 requires TMEM41B for infection is supported by our recent coronavirus genome-wide CRISPR screening and validation results (see accompanying manuscript: (Schneider et al., 2020)).

### Functional TMEM41B is conserved across mammalian and vector species

There are four reported TMEM41B isoforms in humans, however, only isoform 1 encodes a fully intact VTT domain. To determine if any of the other three isoforms can support flavivirus infection, we cloned and expressed each isoform in TMEM41B KO cells. Secondary structure predictions indicate that the first ∼47 amino acids of TMEM41B are unstructured (Kelley et al., 2015). Therefore, we also generated a deletion mutant of isoform 1 lacking the first 47 amino acids. A diagram of these TMEM41B constructs is shown in **Figure 3A**. We found that only the full length and N terminal truncated isoform 1 proteins were able to fully support YFV and ZIKV infection in TMEM41B KO HAP1 cells; however, isoform 4, which contains half of the VTT domain, partially supported YFV infection (**Figure 3B**). From this we conclude that the TMEM41B VTT domain is required to support flavivirus infection whereas the N terminus is dispensable.

**Figure 3.**
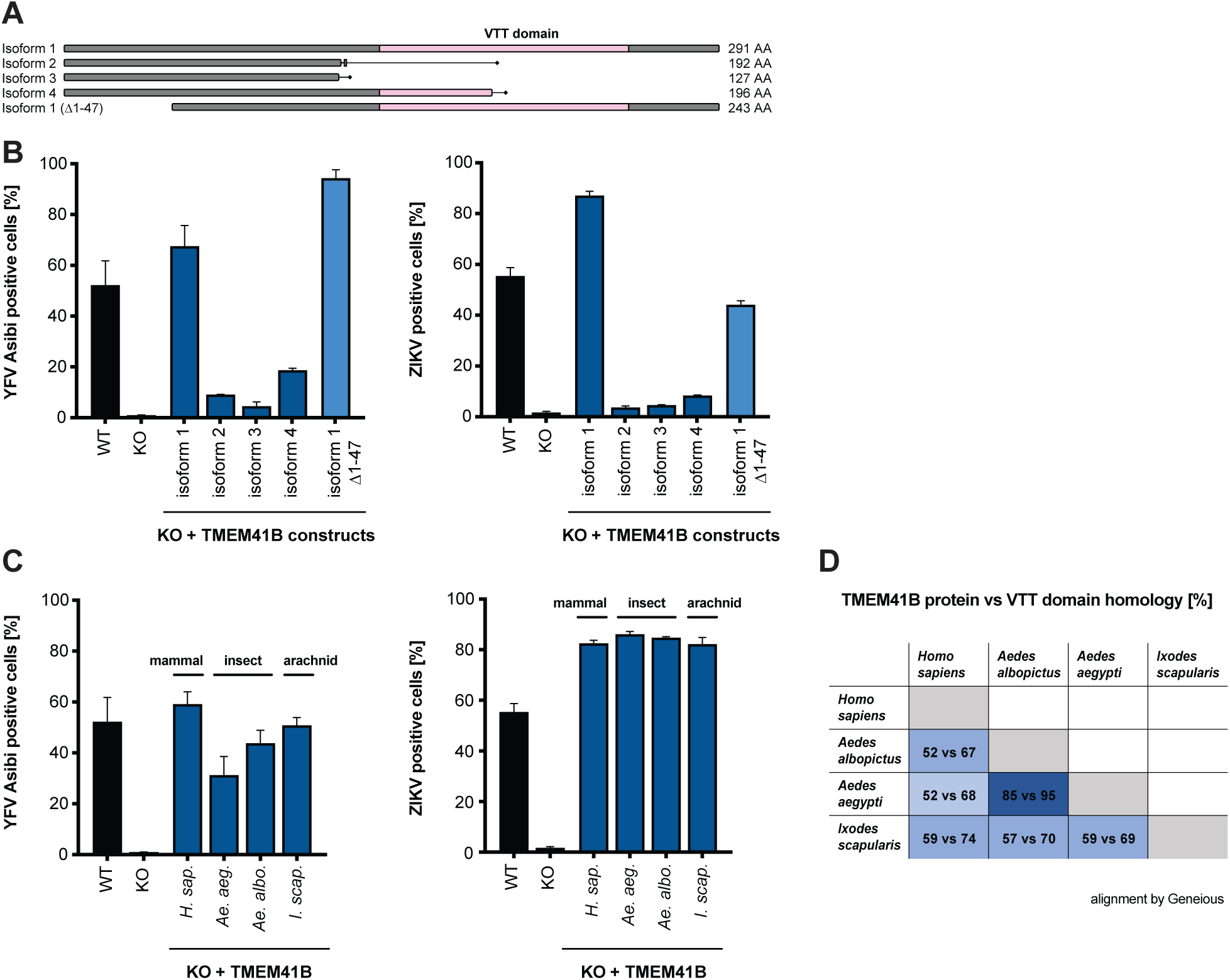
Functional TMEM41B is conserved across mammalian and vector species. **(A)** Graphical representation of human TMEM41B isoforms and deletion construct. VTT domain is shaded pink. Amino acid length of each isoform is indicated. Isoform 1, UniProt ID: Q5BJD5-1; isoform 2, UniProt ID: Q5BJD5-2; isoform 3, UniProt ID: Q5BJD5-3; isoform 4, UniProt ID: E9PJ42. **(B)** WT, TMEM41B KO, and TMEM41B KO cells expressing human TMEM41B isoforms or **(C)** mosquito and tick TMEM41B isoforms infected with YFV Asibi, MOI = 0.1 PFU/cell for 72 h, or ZIKV, MOI = 0.25 PFU/cell for 48 h. **(D)** Table comparing homology between human TMEM41B (isoform 1) and mosquito and tick TMEM41B homologues. Numbers indicate percent amino acids identity for the full-length protein (first number) vs VTT domain only (second number). Protein sequences were aligned using Geneious Software (Geneious 8.1.9. https://www.geneious.com). Cells were analyzed by flow cytometry and plotted as a percentage of viral antigen positive cells. Error bars depict SD for three replicates.

It was previously reported that TMEM41B physically interacts with VMP1, raising the possibility that this interaction may be important for membrane remodeling in the early stages of autophagy (Morita et al., 2019). Based on this observation we hypothesized that evolutionarily divergent TMEM41B homologues may not support flavivirus infection in mammalian cells, given that there are extensive amino acid differences among the mosquito, tick, and human TMEM41B homologues that may prohibit their interaction with human VMP1 (Ishii and Akira, 2005) (**Figure 3D**). To test if mosquito or tick TMEM41B homologues could support flavivirus infection, we expressed TMEM41B homologues from *Aedes aegypti* mosquitoes, *Aedes albopictus* mosquitoes, and *Ixodes scapularis* ticks in TMEM41B KO HAP1 cells and challenged them with YFV and ZIKV. To our surprise, we found that all three TMEM41B homologues supported virus infection in mammalian cells (**Figure 3C**). This suggests that despite amino acid differences, evolutionarily divergent TMEM41B homologues can either functionally interact with human VMP1, or that a direct protein-protein interaction between TMEM41B and VMP1 is not required to support flavivirus infection. As expected, neither TMEM41B KO nor expression of any of these TMEM41B constructs affected hPIV-3 infection (**Figure S3A, B**).

### Naturally occurring TMEM41B SNPs negatively impact flavivirus infection

Viruses and other pathogens have shaped human genetics over millennia. In the face of strong evolutionary pressure, naturally occurring single nucleotide polymorphisms (SNPs) can be enriched or purged from human populations. We investigated if any missense variants existed in human *TMEM41B* and found a SNP (rs78813996) at amino acid position 266 that results in an Ile266Val/Leu substitution with Val being the most prevalent variant. This SNP is nearly absent in European and African populations but is present at ∼3.5% frequency in the Latin American population and at a striking ∼20% frequency in the East and South East Asian population (https://www.ncbi.nlm.nih.gov/snp/rs78813996) (**Figure 4A**).

**Figure 4.**
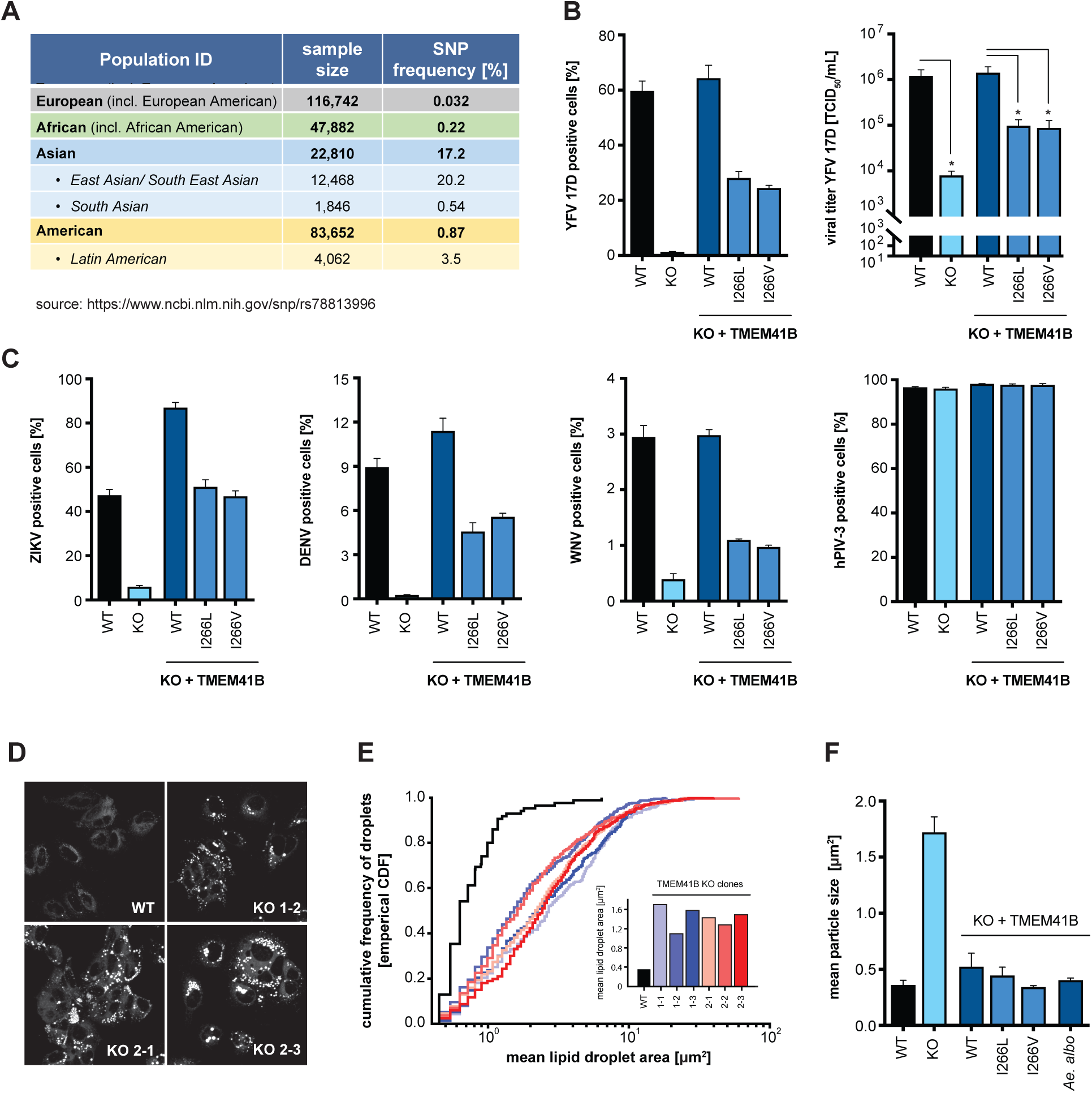
Naturally occurring TMEM41B SNPs negatively impact flavivirus infection but are able to maintain normal lipid distribution in cells. **(A)** Table shows the frequency of a SNP (rs78813996) in *TMEM41B* in several human populations. (B) WT, TMEM41B KO, and TMEM41B KO HAP1 cells expressing WT or TMEM41B SNP variants infected with YFV 17D, MOI = 0.005 PFU/cell for 48 h. Left, cells were analyzed by flow cytometry and plotted as a percentage of viral antigen positive cells. Right, supernatants were collected and titrated by tissue culture infectious dose (TCID50/ml) assay on Huh-7.5 cells. **(C)** Same as panel (B, left) with ZIKV at MOI = 0.25 PFU/cell for 48 h; DENV-GFP at MOI = 0.1 PFU/cell for 96 h; WNV-GFP at MOI = 1 PFU/cell for 72 h; and hPIV-3-GFP at MOI = 0.02 IU/cell for 48 h. **(D)** WT and TMEM41B KO Huh-7.5 cell clones stained with Nile red to visualize lipid droplets. **(E)** Cumulative frequency of droplets plotted vs. droplet area (µm^2^) for six independent single cell clones generated with two independent sgRNAs. Inset shows the mean lipid droplet area (µm^2^) for the six TMEM41B KO clones compared to WT Huh-7.5 cells. **(F)** Mean lipid droplet area (µm^2^) for WT, TMEM41B KO, and TMEM41B KO Huh-7.5 cells (clone 1-1) expressing the indicated TMEM41B variants. Cells were analyzed by flow cytometry and plotted as a percentage of viral antigen positive cells. Error bars depict SD for three replicates.

We engineered these amino acid variants into a TMEM41B expression construct with a red fluorescent protein (tagRFP) fused to the N terminus. Western blot confirmed that both the tagRFP-TMEM41B wildtype and SNP I266L and I266V variants were expressed to similar levels in TMEM41B KO HAP1 cells (**Figure S4A, B**). We then challenged these cells with YFV and found that at 48 h post infection, the percent infection in cells expressing the I266L and I266V variants was reduced by 50% and that these cells produced 10-fold less virus as compared to cells expressing wild type TMEM41B (**Figure 4B**). Consistent with these results, we then tested ZIKV, DENV, and WNV and found similar results to that of YFV, whereas the hPIV-3 control was unaffected (**Figure 4C**). These results suggest that naturally occurring variants in *TMEM41B* reduce infection in cell culture and could potentially affect flavivirus pathogenesis in human populations.

### TMEM41B SNP constructs are able to maintain normal lipid distribution in cells

While culturing TMEM41B KO Huh-7.5 cells, we observed a striking visual accumulation of lipids. To confirm these were indeed lipid droplets, we stained wildtype and six independent TMEM41B KO Huh-7.5 clones with Nile red and found that in all clones lipid droplets are considerably larger as compared to wildtype cells (**Figure 4D**). We then quantified this observation by measuring the area of individual droplets from a single imaging plane in at least 23 cells per clone and found that the average size of individual lipid droplets was up to 10 times larger among TMEM41B KO clones relative to wildtype cells (**Figure 4E**). Moretti et al. reported a similar observation in TMEM41B KO H4 neuroglioma cells and it was later shown that these droplets contain neutral and sterol lipids (Kang et al., 2020; Moretti et al., 2018).

Having found that two naturally occurring TMEM41B variants, I266L and I266V, have a reduced capacity to support flavivirus infection, we next tested if these variants were able to rescue the enlarged lipid droplet phenotype. In addition to the two human variants, we also included the *Aedes aegypti* TMEM41B homologue. As shown in **Figure 4F**, we found that expression of wildtype human and mosquito TMEM41B as well as the two human TMEM41B SNP variants all reduced lipid droplets back to baseline. These results suggest that while these two SNPs negatively impact flavivirus infection, they appear to be fully functional in their ability to maintain normal lipid distribution as measured in this assay.

### TMEM41B colocalizes with flavivirus NS4A and NS4B proteins

Next, we assayed whether the cellular localization of TMEM41B changed upon virus infection. We utilized the tagRFP-TMEM41B expression construct to monitor TMEM41B cellular localization prior to and during flavivirus infection. Previous imaging and proteomics experiments have shown that TMEM41B localizes to ER membranes (Moretti et al., 2018; Morita et al., 2019). Our observations in TMEM41B KO HAP1 cells expressing tagRFP-TMEM41B are consistent with ER localization. Interestingly, however, upon infection with either YFV or ZIKV, we observed a striking re-localization of tagRFP-TMEM41B from a diffuse reticular-like pattern to a large cytosolic aggregate that co-localizes with viral nonstructural proteins, NS4A (ZIKV) and NS4B (YFV) (**Figure 5A**). NS4A and NS4B are multipass transmembrane proteins known to induce membrane curvature, which, together with additional nonstructural viral and likely host proteins, form the viral RNA replication complex.

**Figure 5.**
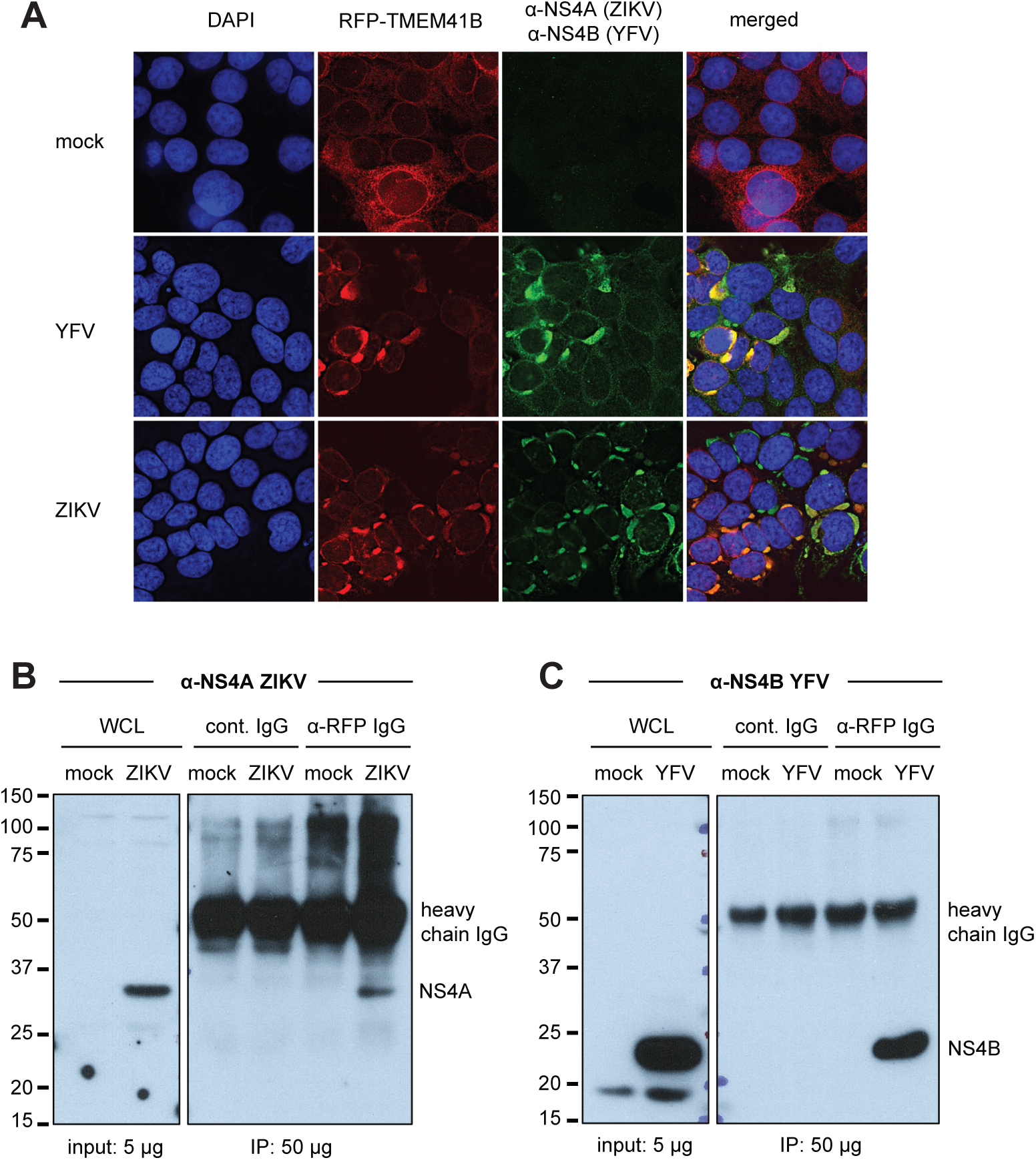
TMEM41B colocalizes with flavivirus NS4A and NS4B proteins. **(A)** TMEM41B KO HAP1 cells expressing RFP-TMEM41B visualized in uninfected cells (mock) and YFV- and ZIKV-infected cells for 24 h at MOI = 1 and 2.5 PFU/cell, respectively). Anti-NS4A (ZIKV) and anti-NS4B antibodies detect viral antigens. Yellow-orange color in the merged column shows the colocalization of TMEM41B with viral antigens. DAPI stains cell nuclei. **(B)** Western blot shows that an anti-RFP antibody which recognizes RFP-TMEM41B (but not an IgG antibody control) co-immunoprecipitates ZIKV NS4A in HAP1 cells (MOI = 2.5 PFU/cell for 24 h). **(C)** Same as (B) but with YFV infection (MOI = 1 PFU/cell for 24 h) and visualized with an antibody that detects YFV NS4B. WCL = whole cell lysate. Of note, the heavy chain of the capture antibodies is detected by the secondary antibody protein A-HRP as indicated ∼ 52 kDa. See also Figure S5.

To further confirm that TMEM41B co-localizes with NS4A and NS4B, we next performed co-immunoprecipitation (co-IP) experiments using an anti-RFP antibody to immunoprecipitate tagRFP-TMEM41B. We found that NS4A in lysates from ZIKV-infected cells and NS4B in lysates from YFV-infected cells co-immunoprecipitated with tagRFP-TMEM41B, whereas no NS4A or NS4B signal was detected in immunoprecipitates from lysates from uninfected cells or infected cell lysates immunoprecipitated with an IgG isotype control antibody (**Figure 5B, C**). We probed these blots for β-actin to confirm that similar amounts of lysate were used as input and we probed for RFP to confirm that equal amounts of tagRFP-TMEM41B were immunoprecipitated in both uninfected and infected cells (**Figure S5A, B**). All together, these results suggest that TMEM41B is likely recruited to sites of viral RNA replication either indirectly or potentially by direct interaction with NS4A and/or NS4B.

### Adaptive mutations in NS4A and NS4B restore flavivirus infectivity in TMEM41B KO cells

Viruses exist as diverse populations of variants and can readily adapt to new environments. These genetic adaptations can highlight viral proteins or viral RNA structures that are under selective pressure in a given context. Throughout our experiments, we often observed rare YFV and ZIKV antigen positive cells after infecting clonal TMEM41B KO populations. To gain potential insight into which stage of the virus replication cycle was affected by the absence of TMEM41B, we sought to capitalize on these rare infection events to select for adaptive mutations within the viral genome. We did this by serial passaging of ZIKV-infected cells and culture supernatants to select viral variants capable of replicating in TMEM41B-deficient cells. We performed virus supernatant passaging experiments in human TMEM41B KO A549 and HAP1 cells, and in parallel, we passaged ZIKV-infected TMEM41B KO mosquito Aag2 cells. Serial passages were performed in multiple independent lineages for each cell type, and we similarly passaged virus in wildtype cells to control for cell culture adaptive mutations that arise which are unrelated to TMEM41B KO. **Figure 6A** shows a graphical outline of these experiments.

**Figure 6.**
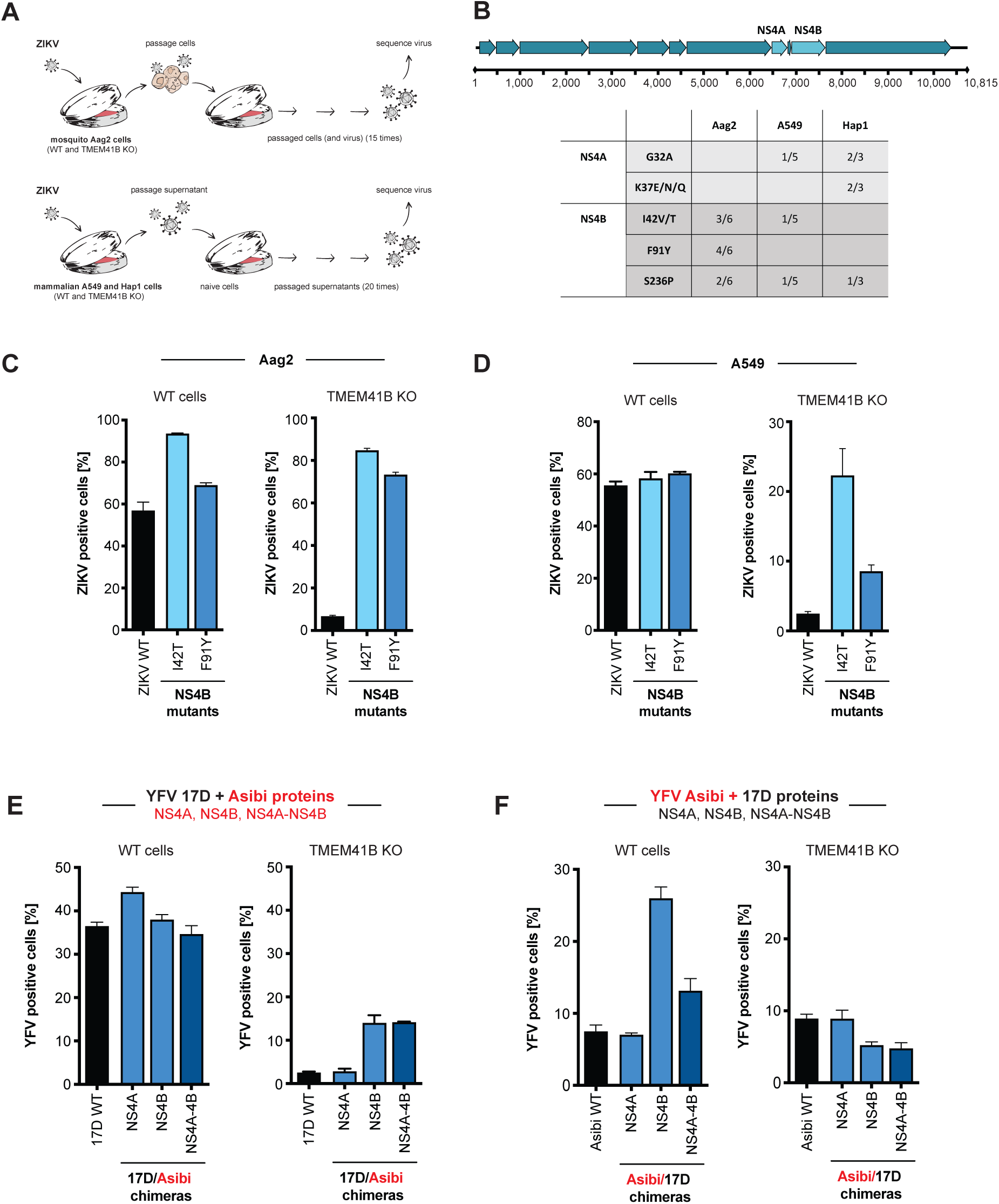
NS4A and NS4B mutations bypass TMEM41B deficiency. **(A)** Graphical schematic of virus adaptation experiment. Top, WT Aag2 and two TMEM41B KO clones were infected with ZIKV (MOI = 0.5 PFU/cell) in triplicate. Cells were passaged every 2-3 days for 15 passages after which supernatants were collected and viral RNA was reverse transcribed and sequenced with next generation sequencing (NGS). Bottom, A549 WT cells together with two TMEM41B KO clones, and HAP1 WT cells together with one TMEM41B KO clone, were infected with ZIKV (MOI = 5 PFU/cell) in triplicate. Supernatants were diluted 1:2 and used to inoculate naïve cells every 3-4 days for 20 passages after which supernatants were collected and viral RNA was reverse transcribed and sequenced with next generation sequencing (NGS). **(B)** Summary of ZIKV adaptive mutations. Top, graphical representation of the ZIKV genome with mature (proteolytically processed) viral proteins shown as arrows and nucleotide positions listed below. NS4A and NS4B, are labeled and colored in light blue. Bottom, table of the amino acids where missense mutations were identified after passaging. Only mutations that were present in at least three independent passages (irrespective of cell type) and that did not appear in virus populations passaged in WT cells are shown. Boxes show the number of replicates a given mutation was identified over the total number of replicates analyzed for that cell type. Of note, out of six initial replicates for the A549 TMEM41B KO clones, only five were able to be carried over the course of 20 passages. **(C)** Missense mutations I42T and F91Y were independently engineered in a ZIKV infectious clone and used to generate virus stocks to infect WT (left) or TMEM41B KO (right) Aag2 cells (MOI = 0.001 PFU/cell for 72 h). The virus stocks used for infection are listed below the bars. **(D)** Same experiment as in (C) in WT and TMEM41B KO A549 cells (MOI = 0.05 PFU/cell for 48 h). **(E-F)** YFV 17D and Asibi chimeric viruses were generated and used to infect WT and TMEM41B KO A549 cells. (E) 17D backbone with Asibi NS4A and NS4B proteins (MOI = 0.1 PFU/cell for 48 h). (F) Asibi backbone with 17D NS4A and NS4B proteins (MOI = 0.1 PFU/cell for 48 h). Cells were analyzed by flow cytometry and plotted as a percentage of viral antigen positive cells. Error bars depict SD for three replicates.

After 15-20 rounds of passaging, we deep-sequenced viral cDNA from cell supernatants to identify viral variants that were enriched in virus populations passaged in TMEM41B KO cells relative to virus populations passaged in wildtype cells. To qualify as a potential adaptive mutation, we applied a stringency filter that required mutations to be present in at least three independent passages (regardless of cell type) and *not* present in virus that had been passaged in wildtype cells. Remarkably, all mutations that met these criteria were located in NS4A and NS4B. Further, NS4B mutations at positions Ile42 and Ser236 were selected in both mosquito and mammalian cells (**Figure 6B**).

To confirm that these mutations we selected enable ZIKV to infect TMEM41B-deficient cells, we chose to introduce two mutations in NS4B, Ile42Thr and Phe91Tyr, into the infectious cDNA clone of the Cambodian ZIKV strain FSS13025 (Shan et al., 2016), and we generated viral stocks. **Figure 6C and D** demonstrates that these individual point mutations in NS4B render ZIKV infectious in both mosquito (Aag2) and mammalian (A549) TMEM41B KO cells.

### Wild and vaccine strain chimeras overcome TMEM41B deficiency

As shown in **Figure 2E** and **S2A**, we observed that two highly similar strains of YFV— the parental Asibi strain and the 17D vaccine strain—were differentially sensitive to TMEM41B deficiency in A549 cells. The live-attenuated YFV 17D strain differs from the Asibi strain by only 31 amino acids, three of which are located within NS4A (V107I) and NS4B (M95I and Y232H) (dos Santos et al., 1995). Given the strain-specific differences we observed in TMEM41B KO A549 cells, together with the observation that single amino acid changes in NS4A and NS4B could allow flavivirus replication in TMEM41B KO cells, we next generated 17D:Asibi chimeric viruses by swapping NS4A and NS4B individually, or by swapping the region containing both NS4A and NS4B which includes the 2K protein. Remarkably, we found that in the 17D viral background, NS4B from the Asibi strain significantly increased infection in TMEM41B KO A549 cells, whereas the NS4B from the 17D strain slightly decreased the infectivity of the Asibi strain in TMEM41B KO cells. In contrast, the NS4A changes had no impact on infection. Importantly, parallel infections in wildtype A549 cells show that the results observed in the KO cells are not due to an overall increase or decrease in viral replication, but rather were likely due to a differential requirement for TMEM41B in this cellular context (**Figure 6E, F**).

### TMEM41B KO cells mount an exaggerated innate immune response to flavivirus inoculum

During the process of studying infection of TMEM41B KO cells, we observed an apparent cytotoxicity with high virus inocula (MOI > 5 PFU/cell) that was not seen with similar inocula to wildtype cells. We hypothesized that this cytotoxicity may have resulted from an exaggerated innate immune response due to increased exposure of viral double-stranded RNA (dsRNA) in TMEM41B-deficient cells. This hypothesis is based on recent evidence supporting a role for TMEM41B in mobilizing lipids and inducing membrane curvature (Moretti et al., 2018; Morita et al., 2019), and our own data which indicates that TMEM41B is likely recruited to flavivirus replication complexes. Flavivirus replication complexes are sites at which viral RNA replication machinery assembles within invaginations into the ER lumen utilizing host curvature-stabilizing proteins (Aktepe and Mackenzie, 2018; Neufeldt et al., 2018; Rajah et al., 2020). Within these structures, dsRNA intermediates that form during viral RNA replication are protected from host innate immune sensors that survey the cytosol for pathogen associated molecular patterns (PAMPs). In the absence of TMEM41B, viral RNA replication may initiate in aberrant replication complexes that are exposed to the cytosol.

To test if the observed cytotoxicity was due to innate immune activation, we inoculated wildtype and TMEM41B KO HAP1 cells with a high YFV 17D virus inoculum (MOI = 0.4 PFU/cell), then quantified viral RNA and mRNA encoding 2’-5’-oligoadenylate synthetase 1 (OAS1) by qPCR at 24 h post infection. OAS1 is a classic interferon (IFN) stimulated gene (ISG) induced by type I and III IFNs. As shown in **Figure 7A**, we found that YFV 17D RNA levels were 100-fold lower in TMEM41B KO cells compared to wildtype cells, consistent with an inability to replicate. Supporting our hypothesis, OAS1 was highly induced in the KO cells but not induced in wildtype cells. To determine if OAS1 induction was due to the virus itself or functional viral RNA, we inoculated both wildtype and TMEM41B KO cells with viral particles inactivated by UV light. UV-inactivated virus failed to induce OAS1 expression in both cell lines suggesting that viral protein production and perhaps initiation of viral RNA replication was required to induce innate immunity (**Figure 7B**). To control for the possibility that TMEM41B KO cells are hypersensitive to PAMPs in general, we infected both wildtype and TMEM41B KO cells with a recombinant influenza A virus (IAV) lacking a critical innate immune antagonist protein, NS1 (IAV ΔNS1). This virus is well known for its defect in antagonizing IFN production and ISG expression (Garcia-Sastre et al., 1998; Wang et al., 2000). We found that both wildtype and TMEM41B KO responded to IAV ΔNS1 as expected, with wildtype cells expressing even slightly higher amounts of OAS1 mRNA than TMEM41B KO cells (**Figure 7B**).

**Figure 7.**
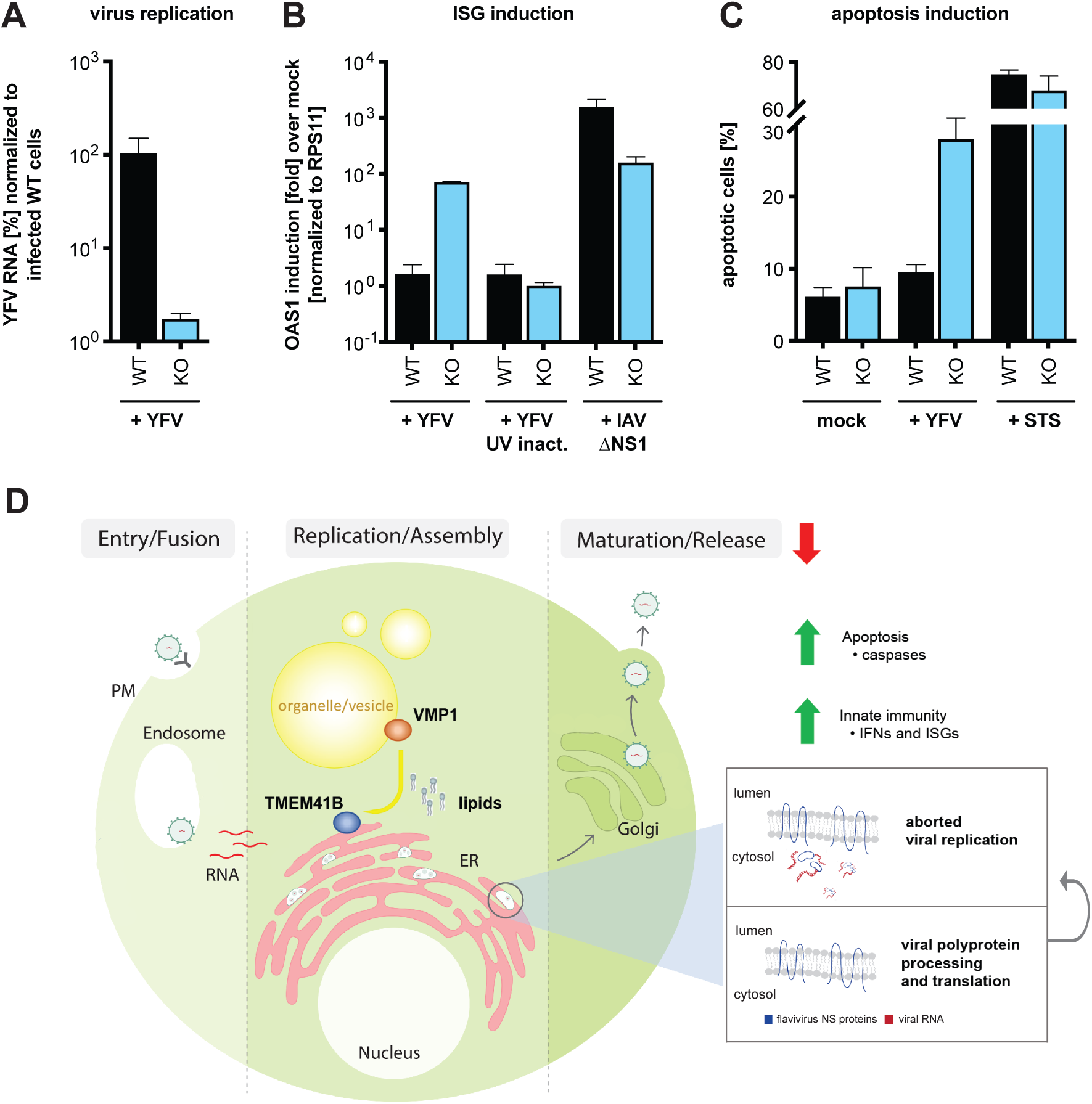
TMEM41B KO cells mount an exaggerated innate immune response to flavivirus inoculum. **(A)** Viral RNA quantified by qRT-PCR from WT and TMEM41B KO HAP1 cells infected with YFV 17D MOI = 0.4 PFU/cell for 24 h. **(B)** OAS1 mRNA quantified by q-RT-PCR from WT and TMEM41B KO HAP1 cells infected with YFV 17D (with and without UV-inactivation) MOI = 0.4 PFU/cell for 24 h or with IAV (ΔNS1) MOI = 0.1 PFU/cell for 24 h. **(C)** WT and TMEM41B KO HAP1 cells untreated (mock), infected with YFV 17D MOI = 4 PFU/cell for 24 h, or treated with 250 nM staurosporine (STS) and assayed to detect apoptotic cells via Annexin-V staining. Cells were analyzed by flow cytometry and plotted as a percentage of viral antigen positive cells. Error bars depict standard deviation (SD) of n = three replicates. **(D)** Model for the role of TMEM41B in the flavivirus life cycle. The model is described in the main text.

Upon detection of viral PAMPs, cells secrete IFN to protect neighboring cells from infection then subsequently initiate apoptosis (Drappier and Michiels, 2015; Gusho et al., 2020; Schwartz and Conn, 2019). To assay if the cytotoxicity we observed in TMEM41B KO cells after inoculation with high viral loads was in fact apoptosis, we inoculated wildtype and TMEM41B KO cells with a high YFV 17D inoculum (MOI = 4 PFU/cell) and quantified the number of apoptotic cells at 24 h post inoculation. As shown in **Figure 7C**, we observed an approximate three-fold increase in the percentage of apoptotic cells in TMEM41B KO cells compared to wildtype cells. Importantly, at this early time point, there was almost no increase in the percentage of apoptotic cells in the infected wildtype cell population as compared to mock-infected cultures. Further, the positive control, staurosporine (STS), induced apoptosis in approximately 70% of the cells, with no notable difference between wildtype and TMEM41B KO cells (**Figure 7C**).

## DISCUSSION

Genome-wide CRISPR-Cas9 gene disruption screens are a powerful method of discovery. We applied this method to discover cellular host factors that flaviviruses require to replicate in human cells. As a testament to the robustness of this approach, our results along with CRISPR-Cas9 screens performed by research groups using different flaviviruses, different cell types, and different sgRNA libraries identified many shared genes and biological pathways. Screens, however, are often only the first step towards understanding new virus-host biology.

From our loss-of-function screens using a ZIKV isolate from Puerto Rico (PRVABC59) and the Asibi strain of YFV, we noted a strong dependence on an obscure host factor with a generic name: TMEM41B. During the course of our studies, TMEM41B was independently identified by several groups that performed CRISPR-Cas9 loss-of-function screens designed to uncover novel regulators of autophagy. As a result, a cellular role for TMEM41B began to emerge.

In our validation studies we found that all flaviviruses we tested, including both mosquito-borne and tick-borne flaviviruses, required TMEM41B for infection and replication. Further, we found that TMEM41B was required in the mosquito vector as well, supporting a conserved role for TMEM41B in flavivirus infection in distantly related species. Given that TMEM41B is required for flavivirus infection and that TMEM41B is also required for autophagy, the question arose: is autophagy required for flavivirus infection? To address this question, we generated CRISPR KO cell lines lacking *BECN1, ATG5*, and *ATG7* since these genes are known to have critical roles in cellular autophagy. Our results suggest that under the conditions we tested the full autophagy pathway is not essential for flavivirus infection. This is also indirectly supported by the fact that *VMP1* and *TMEM41B* are the only hits shared among our ZIKV CRISPR-Cas9 screen and a CRISPR-Cas9 autophagy screen. This led us to hypothesize that flaviviruses may hijack TMEM41B and VMP1 for their ability to remodel host cell membranes–a feature required both for autophagy and for the formation of viral replication complexes.

Although every flavivirus we tested required TMEM41B in most cell types, there were several virus-cell combinations we tested where TMEM41B was either not required or played a less essential role in virus infection. This conundrum may be explained by cell type-specific differences in the abundance of structurally or functionally related cellular host proteins such as VMP1 and TMEM41A, which we found can partially support flavivirus infection in the absence of TMEM41B. These findings suggest that TMEM41B may be an important cellular determinant that defines flavivirus tissue tropism and pathogenesis. Along these lines, it is notable that a SNP in *TMEM41B* present in approximately 20% of East and South East Asian and 3.5% of Latin American populations has a reduced capacity to support flavivirus infection. While it is nearly impossible to know for sure what drove selection of this SNP in these populations, it is conceivable that a pathogenic flavivirus may have been involved.

While variations in the host environment can influence virus tropism, so too can variations in the virus. Our studies uncovered that two highly related yellow fever viruses, the virulent Asibi strain and the live attenuated 17D vaccine strain, displayed a differential ability to infect JEG3 and A549 cells lacking TMEM41B. The fact that the YFV Asibi and 17D strains differ by only 31 amino acids provided us with an opportunity to potentially isolate the flavivirus TMEM41B dependency down to the level of an individual viral protein. Concurrently, we made additional observations that suggested amino acid differences in NS4A and NS4B between these two strains may be responsible for their differential TMEM41B requirement. These observations included our findings that (i) TMEM41B relocalizes upon flavivirus infection and colocalizes with NS4A and NS4B, (ii) TMEM41B co-immunoprecipitated with NS4A and NS4B in flavivirus-infected cells, and (iii) mutations within ZIKV NS4A and NS4B can compensate for the lack of TMEM41B. Based on these data, we made 17D:Asibi chimeric viruses and found that two amino acid differences within NS4B were sufficient to confer or bypass TMEM41B dependence. Future studies are needed to better understand how only one or a few amino acids changes in NS4A and NS4B are able to bypass the strong TMEM41B requirement during flavivirus infection. Together these results highlight remarkable plasticity even among "essential" host factors and demonstrate that loss of a critical host factor can be overcome by relatively few genetic changes in the viral genome. This contrasts the notion that resistance barriers may be higher when targeting host factors with antiviral therapeutics as opposed to targeting the virus. When dealing with highly diverse viral populations, multi-target drug cocktails are still likely to be important regardless of whether therapies target host or viral factors.

Our final observation where we demonstrate that TMEM41B cells initiate a strong innate immune response when exposed to a large number of virus particles provides mechanistic insights into the role of TMEM41B in the flavivirus replication cycle. Based on our collective results together with what is known about the role of TMEM41B in lipid mobilization and inducing membrane curvature, we propose a model, shown in **Figure 7D**, where upon flavivirus infection and translation of the viral polyprotein, TMEM41B is recruited to sites on the ER membrane together with NS4A and NS4B where replication complexes are forming (Neufeldt et al., 2018; Paul and Bartenschlager, 2015). This recruitment may be through direct protein:protein interaction or through passive diffusion, where, by mobilizing neutral and sterol lipids, TMEM41B helps lower the local free energy imposed by NS4A- and NS4B-induced membrane curvature. We propose that TMEM41B’s role in facilitating membrane curvature is dependent upon a functional interaction between TMEM41B and VMP1, the latter of which is highly mobile and associates with cell organelles, vesicles and lipid droplets (Tabara and Escalante, 2016; Zhao et al., 2017). In the absence of TMEM41B, flavivirus NS4A and NS4B proteins assemble and viral RNA replication initiates, but because replication complexes are improperly formed, dsRNA intermediates are exposed to pattern recognition receptors (PRRs). At this point, the infectious replication cycle is aborted, either due to an innate immune response or simply because the RNA genome cannot be efficiently copied. Future work is needed to thoroughly test this model and understand how TMEM41B works together with viral proteins and other host factors during assembly of flavivirus replication complexes.

Lastly, during our studies, we uncovered a role for TMEM41B in the coronavirus replication cycle. Recently, we and others also identified TMEM41B as a top scoring hit in genome-wide CRISPR-Cas9 loss of function screens designed to identify coronavirus host dependency factors ((Baggen et al., 2020; Wang et al., 2020) and accompanying manuscript: (Schneider et al., 2020)). It will be fascinating to learn if TMEM41B is also required to assemble coronavirus replication complexes. In conclusion, TMEM41B may be a candidate target to inhibit the replication of a broad range of emerging and re-emerging flavivirus and coronavirus pathogens.

## ACKNOWLEDGEMENTS

We thank the following investigators for contributing viral molecular clones and viral stocks: P. Hearing (replication-competent AdV5-GFP), I. Mohr (HSV-1-GFP), P.L. Collins (hPIV-3-GFP), J.K. Rose (VSV-GFP), J.C. de la Torre (LCMV-GFP), P. Palese (IAV), A. Garcia-Sastre (IAV ΔNS1), J.L Whitton (CVB-3-GFP), P. Traktman (VacV-GFP), I. Frolov (WNV-GFP), S. Higgs (CHIKV-GFP), R. Elliott (BUNV-GFP), P.Y. Shi (ZIKV: FSS13025), Centers for Disease Control and Prevention (ZIKV: PRVABC59), and BEI Resources (SARS-CoV-2).

We thank the Rockefeller University Flow Cytometry and Bio-Imaging Resource Center for their training, advice, organization, and for maintaining exceptional facilities. We thank Joe Luna and Francisco J. Sánchez-Rivera for their boundless enthusiasm and for stepping up in time of crisis to allow this manuscript to be written. We thank Yingpu Yu for engaging discussion and generosity. We thank Alison W. Ashbrook and Lauren C. Aguado for critical feedback on short notice. We thank Aileen O’Connell, Santa Maria Pecoraro Di Vittorio, Glen Santiago, Mary Ellen Castillo, Arnella Webson and Sonia Shirley for outstanding administrative or technical support.

Research reported in this publication was supported by the National Institute Of Allergy and Infectious Diseases of the National Institutes of Health under Award Number R01AI124690 (to C. M. Rice). The content is solely the responsibility of the authors and does not necessarily represent the official views of the National Institutes of Health. The project was also supported in part by NIH grant P30 CA008748 (to C. M. Rudin). F.S. received a MD fellowship from Boehringer Ingelheim Fonds (BIF). This work was further supported by the generosity of the Bawd Foundation and the Robertson Foundation.

## AUTHOR CONTRIBUTIONS

Conceptualization: HHH, WMS, KRG, LAM, FS, MRM, JTP, CMR

Methodology: HHH, WMS, KRG

Formal analysis: JTP

Investigation: HHH, WMS, KRG, LAM, FS, BR, EJ, XW, SY, LKM

Supervision: MRM, JTP, CMR

Visualization: HHH, WMS, JTP

Writing - original draft: HHH, WMS

Writing – review & editing: HHH, WMS, LAM, FS, BR, MRM, LKM, JTP, CMR

Funding acquisition: HHH, WMS, CMR, MRM, JTP, CMR

## METHODS

### Screen

HAP1 B3GALT6 KO-Cas9 cells were generated by lentiviral transduction of lentiCas9-Blast, a gift from Feng Zhang (Addgene: #52962; http://n2t.net/addgene: 52962; RRID: Addgene_52962), followed by selection and expansion in the presence of 1.5 μg/ml blasticidin. The human GeCKO library (A and B) (Shalem et al., 2014) was a gift from Feng Zhang and obtained through Addgene (cat. #1000000049). The plasmid was expanded and lentivirus was produced as previously described (Shalem et al., 2014). To deliver the GeCKO (A and B) sgRNA library to cells, 2.03 x 10^8^ HAP1-Cas9 cells were transduced at MOI = 0.4 to achieve ∼1,600-fold overrepresentation of each sgRNA. Two days later, media was replaced with fresh media containing 1.5 μg/ml puromycin and cells were expanded for six additional days prior to seeding for ZIKV or YFV infection.

We seeded cells at 3.5 x 10^6^ cells per T175 flask and we seeded 15 flasks per replicate (in triplicate) for each virus infection as well as Mock. The following day, the media was removed and viruses diluted in 10 ml/flask OptiMEM were added to cells. ZIKV was added at MOI = 0.025 PFU/cell and YFV Asibi was added at MOI = 0.5 PFU/cell. After three hours on a plate rocker at 37 °C, plates were moved to 5% CO2 incubators set to 37 °C. Media was changed three days later. Media was changed every 3-4 days to remove dead cells. Mock cells were passaged every 3-4 days and re-seeded at 3.5 x 10^6^ cells/flask to maintain library complexity. Cells were harvested approximately twenty days post infection.

Genomic DNA (gDNA) was isolated via DNeasy Blood & Tissue Kit (Qiagen: 69504) per the manufacturer’s instructions. The library was amplified from gDNA as previously described (Yau and Rana, 2018). PCR products were agarose gel purified pooled, and sequenced on an Illumina NextSeq 500 at the MSKCC Integrated Genomics Operation using standard Nextera sequencing primers and 50 cycles.

### Analysis of CRISPR-Cas9 genetic screen data

FASTQ files were processed and trimmed to retrieve sgRNA target sequences followed by enumeration of sgRNAs in the reference sgRNA library file using MAGeCK (Li et al., 2014). Z-scores were computed using the following approach: for each condition, the log2 fold change with respect to the initial condition was computed. A natural cubic spline with 4 degrees of freedom was fit to each pair of infected and control cells and residuals were extracted. To obtain gene-wise data, the mean residuals for each group of sgRNAs was calculated, a z-score was computed, and a p-value was determined using a 2-sided normal distribution test. P-values were combined across screens using Fisher’s sumlog approach and corrected for multiple testing using the method of Benjamini & Hochberg.

### Cell Culture

HAP1 (WT and B3GALT6 KO) (human; sex: male, chronic myeloid leukemia-derived) cells were obtained from Thijn Brummelkamp (Netherlands Cancer Institute) and cultured in Iscove’s Modified Dulbecco’s Medium (IMDM, Gibco) supplemented to contain 10% fetal bovine serum (FBS, Hyclone, GE Healthcare) and 1% non-essential amino acids (NEAA, ThermoFisher Scientific). A549 cells (human; sex: male, lung epithelial), JEG3 cells (human; sex: female, placenta epithelial), HEK293 derivative Lenti-X™ 293T cells (human; sex: female, kidney epithelial) obtained from Takara (cat. 632180), Huh-7.5 heptoma cells (human; sex: male, liver epithelial) (Blight et al., 2002) and MDBK cells (bovine; sex: male, kidney epithelial) were cultured in Dulbecco’s Modified Eagle Medium (DMEM, Gibco) supplemented to contain 10% FBS (or 10% horse serum [ThermoFisher Scientific] in case of MDBK cells) and 1% NEAA. All mammalian cell lines were obtained from the ATCC unless stated otherwise, cultured at 37 °C and incubated with 5% CO_2_. Mosquito C6/36 (*Ae. albopictus*, sex: unspecified, neonate larva) and Aag2 (*Ae. aegypti*, sex: unspecified, neonate larva) cells were obtained from ATCC and Dr. Maria Carla Saleh, respectively. Cells were cultured in Leibovitz’s L-15 Medium, without phenol red (ThermoFisher Scientific), supplemented to contain 10% FBS, 1% NEAA, and ∼0.3 g/L tryptose phosphate broth (Sigma-Aldrich) at 28 °C in the absence of CO_2_. All cell lines tested negative for contamination with mycoplasma and with the exception for Huh-7.5 cells have not been further authenticated.

### Antibodies and Chemicals

Primary antibodies used for western blot include rabbit anti-TMEM41A (Proteintech: 20768-1-AP; RRID: AB_10693679; 1:1,000), rabbit anti-TMEM41B (Abnova: PAB20785; RRID: AB_10965049; 1:1,000), rabbit anti-VMP1 (Cell Signaling Technology: 12929; RRID: AB_2714018; 1:1,000), rabbit anti-ATG5 (Abcam: ab108327; RRID: AB_2650499; 1:5,000), rabbit anti-ATG7 (Abcam: ab52472; RRID: AB_867756; 1:50,000), rabbit anti-LC3B (Abcam: ab192890; RRID: AB_2827794; 1:2,000), rabbit anti-TagRFP (Evrogen: AB233; RRID: AB_2571743; 1:1,000) and mouse anti-β-actin conjugated to HRP (Sigma-Aldrich: A3854; RRID: AB_262011; 1:50,000).

Primary antibodies used for IF, flow cytometry and IP include mouse anti-YFV (Santa Cruz Biotechnology: sc-58083; RRID: AB_630447; 1:1,000), mouse-anti-flavivirus group antigen (Millipore: MAB10216; RRID: AB_827205; 1:500), pan-flavivirus hyperimmune mouse acites fluid (HMAF, kindly provided by Tom Ksiazek, CDC; 1:200), rabbit anti-YFV NS4B (GeneTex: GTX134030; 1:1,000), rabbit anti-ZIKV NS4A (GeneTex: GTX133704; 1:1,000), rabbit-anti-SARS-CoV-2 N (GeneTex: GTX135357; RRID: AB_2868464; 1:2,000), mouse-anti-IAV nucleoprotein (Millipore: MAB8251; RRID: AB_95293; 1:1,000), anti-BVDV polyclonal antibody B224 (kindly provided by Kenny Brock, Auburn University; 1:500), rabbit anti-TagRFP (see above) and normal rabbit IgG (Cell Signaling Technology: 2729; RRID: AB_1031062).

Secondary antibodies, proteins and staining solutions used for IF, flow cytometry and IP include goat-anti-rabbit IgG conjugated to HRP (ThermoFisher Scientific: 31462; RRID: AB_228338; 1:5,000), goat anti-mouse IgG conjugated to HRP (Jackson ImmunoResearch Labs: 115-035-146; RRID: AB_2307392; 1:10,000), goat anti-bovine IgG conjugated to FITC (ThermoFisher Scientific: A18752; RRID: AB_2535529; 1:1,000), goat anti-mouse IgG conjugated to AF647 (Molecular Probes: A-21235; RRID: AB_2535804; 1:1,000), goat-anti-mouse IgG conjugated to AF488 (ThermoFisher Scientific: A-11001; RRID: AB_2534069; 1:1,000), goat anti-rabbit IgG conjugated to AF488 (Molecular Probes: A-11008; RRID: AB_143165; 1:2,000) and Protein A-HRP (Thermo Fisher Scientific: 10-1023). Hoechst 33342 solution (ThermoFisher Scientific: 62249) was used at 20 μM for nuclear stain. Nile Red Staining Kit (Abcam: ab228553) was used according to the manufacturer’s instructions to stain for intracellular lipid droplets.

Drugs and antibiotics were used at concentrations indicated in figure legends and include chloroquine diphosphate salt (Sigma: C6628), Torin 1 (Selleckchem: S2827), doxycycline hyclate (Sigma: D9891), puromycin dihydrochloride (Sigma: 8833), blasticidin S (Sigma) and polybrene (Sigma: TR-1003-G).

### Lentivirus Production and Transduction

All ORFs were cloned into a modified pTRIPZ vector for lentivirus delivery to cells. All plasmid sequences are available upon request. Lentivirus stocks were generated in Lenti-X™ 293T cells by co-transfection of plasmids expressing (1) the ORF of interest (2), HIV gag-pol, and (3) the vesicular stomatitis virus glycoprotein (VSV-G) in a ratio of 0.55:0.35:0.1 using Lipofectamine^TM^ 2000 at a ratio of 2.5 µl reagent to 1 µg DNA. One day prior to transfection, cells were seeded at 4 x 10^5^ cells per well of poly-L-lysine coated 6-well plates. Cells were transfected the following day and six hours post transfection media was removed and lentivirus was collected overnight in 2 ml per well. Supernatants were collected at 24 and 48 h, filtered using 0.45-micron syringe filters, and stored at -80 °C

For lentiviral transduction, 3 x 10^5^ cells were transferred to 12-well plates in suspension and transduced with several lentivirus dilutions by spinoculation at 1,000 *x g* for 60 min at 37 °C in medium containing 3% FBS, 20 mM HEPES, and 4 µg/ml polybrene. The following day cells were trypsinized and moved to duplicate wells of 6-well plates. At approximately 48 h post transduction one well of each duplicate was selected with 2 µg/ml puromycin. When selection was complete, the selected and unselected replicates were counted to determine approximate MOI and we proceeded to expand cells that were transduced at MOI < 0.3 to ensure that most cells in the population contained a single integrant.

### Generation and Validation of CRISPR KO clones

#### Mammalian cells

Small guide RNAs (sgRNA) for CRISPR editing were designed using crispor.tefor.net (Haeussler et al., 2016) and cloned into lentiCRISPRv2 plasmid CRISPR KO clones were generated by transfection with the relevant guides, selected with 2 µg/ml puromycin for four days prior to single-cell seeding into 96-well plates in the absence of puromycin at a dilution of 0.7 cells/well to obtain single-cell clonal populations. Clones were expanded and screened for protein knockout by western blot analysis or sequencing analysis. Genomic DNA was extracted using the Qiagen DNeasy Blood and Tissue Kit (Qiagen) and used as a template for amplification of an approximately 500-1,000 bp region flanking the PAM site. Discrete bands were gel-extracted and gene disruption was confirmed by submitting samples to the CCIB DNA Core Facility, Massachusetts General Hospital (Cambridge, MA, USA) for high throughput amplicon sequencing.

We used the following sgRNAs in these studies. sgRNAs targeting TMEM41B in HAP1, A549, JEG3, and Huh-7.5 cells: 5’-GTCGCCGAACGATCGCAGTT -3’ and 5’-GCTCACCACACGACCCCCGT -3’. sgRNA targeting TMEM41B in MDBK cells: 5’-ATACTGAGAAATATAGAGCC -3’. sgRNA targeting TMEM64 in HAP1 cells: 5’-GCTCCAGATAGGCGCCGAGC -3’. sgRNA targeting TMEM41A in HAP1 cells: 5’-TCGCGTCGACAGCAAGTACA -3’. sgRNA targeting VMP1 in HAP1 cells: 5’-GCTCTGGCCATGAAATATGG -3’. sgRNA targeting ATG5 in HAP1 cells: 5’-TTCCATGAGTTTCCGATTGA -3’. sgRNA targeting ATG7 in HAP1 cells: 5’-CCTAGCTACATTGCAACCCA -3’. sgRNA targeting BECN1 in HAP1 cells: 5’-CCTGGATGGTGACACGGTCC -3’. All oligos were purchased from IDT.

#### Mosquito cells

CRISPR/Cas9 plasmids were generated from the pDDC6 vector, which encodes the human codon-optimized *Streptococcus pyogenes* Cas9^D10A^ nickase (*hSpCas9D10A*; a gift from Peter Duchek) (Addgene plasmid: #59985; http://n2t.net/addgene: 59985; RRID: Addgene_59985). For expression in mosquito cells, we replaced the *dme* phsp70 promoter (*hsp70Bb*) in pDCC6 with the *Ae. aegypti* polyubiquitin promoter (*aae PUb*) and we replaced the *dme* U6-2 promoter with the *Ae. aegypti* U6 promoter (*aae* U6; AAEL01777419). Plasmid sequence is available upon request. The following sgRNAs were cloned into the abovementioned plasmid. sgRNAs targeting TMEM41B homologue (AAEL022930-RB) in Aag2 cells: 5’-CAAAGATCTCTACTACCTGG-3’ and 5’-TATGTAGACGAGGACCACGC-3’. sgRNAs targeting TMEM41B homologue (AALF002881-RA) in C6/36 cells: 5’-CAAACAGTTGGGACGAGTGC-3’ and 5’-TGTAGCTGAGGAGGTCTTCC-3’. All oligos were purchased from IDT. Cells were transfected with the appropriate plasmids using Fugene HD Transfection Reagent (Promega) according to the manufacturer’s protocol. Cells were seeded at ∼50% confluency and complexes were formed using a ratio of 3:1 transfection reagent to plasmid DNA. To generate single cell clones, cells were seeded in 96-well plates (∼0.7 cells/well) in 50% fresh media, and 50% conditioned media and expanded. Gene disruption was confirmed by high throughput amplicon sequencing and described above.

### Virus Production and Infections

The generation of viral stocks for additional viruses tested have been previously described: DENV-GFP (Schoggins et al., 2012) (derived from IC30P-A, a full-length infectious clone of strain 16681), WNV-GFP (McGee et al., 2010) (derived from pBELO-WNV-GFP-RZ ic), HCV-RFP (Liu et al., 2015) (based on Jc1-378-1), BVDV-NADL (Mendez et al., 1998), CHIKV-GFP (Tsetsarkin et al., 2006) (derived from pCHIKV-LR 5′GFP), hPIV3-GFP (Zhang et al., 2005) (based on strain JS), BUNV-GFP (Shi et al., 2010) (based on rBUN-del7GFP), VacV-GFP (Schoggins et al., 2014) (based on strain Western Reserve), HSV-1 US11-GFP (Benboudjema et al., 2003), replication-competent AdV5-EGFP (generously provided by Patrick Hearing, Stony Brook University), VSV-GFP (Dalton and Rose, 2001), LCMV-GFP (generously provided by J.C. De la Torre, Scripps Research), influenza viruses A/WSN/33 and A/PR/8/34 ΔNS1 (generously provided by Peter Palese and Adolfo Garcia Sastre, Mount Sinai School of Medicine). Lenti-GFP was generated in Lenti-X™ 293T as described above. CVB3-GFP (Feuer et al., 2002) (derived from infectious clone pMKS1-GFP) was amplified in HeLa cells and titrated by TCID_50_. SARS-CoV-2 (WA1/2020 obtained from BEI Resources) and ZIKV (PRVABC59 obtained from the CDC, Ft. Collins) were amplified in Huh-7.5 and HAP1 cells respectively, and titrated by standard plaque assay on Huh-7.5 cells. YF viruses (17D and Asibi) and the Cambodian ZIKV (Shan et al., 2016) (strain: FSS13025, generously provided by Pei-Yong Shi) were generated from infectious clones as described below. Experiments with YFV (strain: Asibi), WNV-GFP, CHIKV-GFP and SARS-CoV-2 were carried out in biosafety level 3 (BSL3) containment in compliance with institutional and federal guidelines. The tick-borne viruses POWV (strain: Byers, VSPB 812027), TBEV (strain: Hypr, VSPB 812222 and Sofjin, VSPB 812223), OHFV (strain: Bogoluvovska, VSPB 812005), KFDV (strain: P9605, VSPB 811996) and AHFV (strain: 200300001, VSPB 200300001) were generated at the CDC and experiments were carried out in BSL4 containment in compliance with institutional and federal guidelines (Flint et al., 2014; Zaki, 1997).

### Generation of recombinant YFV and ZIKV

Recombinant viruses for the Cambodian ZIKV (strain: FSS13025), the YFV vaccine strain 17D and wildtype Asibi strain, as well as the YF chimeric 17D and Asibi viruses were generated using plasmids encoding the infectious clones. To generate mutant and chimeric viruses, overlapping oligos were created to either introduce point mutations or to allow the amplification of specific coding sequences. PCR reactions were run under standard conditions using KOD Hot Start Master Mix (Sigma-Aldrich: 71842). After subsequent ligation with T4 ligase (NEB: M0202), the plasmid DNA was amplified in competent DH5-ɑ E.*coli* cells. All plasmids encoding WT and mutant/chimeric viruses were sequenced for the absence of unwanted mutations within the viral coding region. For *in vitro* transcription, 2 µg linearized plasmid DNA was used in a 30 µl reaction with the mMessage mMachine SP6 Transcription Kit (Invitrogen: AM1340). RNA integrity was examined on a 1% agarose gel. Huh-7.5 cells were subsequently electroporated with 4 µg of YFV RNA mixed into 6 x 10^6^ cells, while C6/36 cells were used for electroporating ZIKV RNA under similar conditions. Electroporated cells were incubated for 48-72 h before collecting and titrating supernatants by standard plaque assay in Huh-7.5 cells.

All virus infections and quantification were performed using the doses and timepoints reported in the figure legends for quantification by flow cytometry or microscopy. For infections, cells were seeded at a density of 1 x 10^4^ cells/well into black, glass-bottom 96-well plates (PerkinElmer: 6055302) and at 2.5 x 10^4^ - 5 x 10^4^ cells/well into regular tissue culture 24-well plates. The next day, cells were washed 1x with Opti-MEM (Gibco) and adsorbed with virus inoculum (50 µl for 96-well and 200 µl for 24-well plates) prepared in Opti-MEM at 37 °C. After 1-2 h inoculum was removed, cells were washed with Opti-MEM, fresh media was added, and cells were incubated at 37 °C. At the final time point dependent on viral replication kinetics, cells in 24-well plates were lifted with Accumax cell dissociation medium (Sigma: A7089) and fixed with 2% paraformaldehyde (PFA). Cells infected with GFP/RFP reporter viruses were pelleted at 930 *x g* for 5 min at 4 °C, resuspended in cold phosphate-buffered saline (PBS) containing 3% FBS and stored at 4°C until analysis using a LSRII flow cytometer (BD Biosciences). FlowJo software v10 (Treestar) was used to obtain the percentage of GFP/RFP-expressing cells, which were graphed using Prism8 (Graphpad).

For non-fluorescent reporter viruses, cells were washed 1x in PBS (+ 2% FBS), permeabilized in 1x BD perm (BD Biosciences: 554723) for 1 h at RT followed by overnight incubation with primary antibody. Cells were then washed three times with BD perm, incubated with secondary antibody for 1 h at RT, washed an additional three times with BD perm, and once with PBS (+ 2% FBS) before being transferred to FACS tubes for flow-cytometry analysis similar as for aforementioned reporter viruses.

Cells infected in 96-well plates were fixed with 10% neutral-buffered formalin, permeabilized with 0.1% Triton X-100 in PBS for 10 min at RT, washed in PBS with 0.01% Tween20 (PBS-T) and incubated with primary antibody (pan-flavivirus HMAF) overnight. The secondary antibody was used at RT for 1 h together with NucBlue (ThermoFisher) to stain for nuclei. Plates were subsequently read using the Perkin Elmer Operetta confocal high content imager with a 10x objective. Images were analyzed with a cell segmentation workflow using Harmony software v4.1 Perkin Elmer, Waltham, MA.

### Immunofluorescence

Unless otherwise described, cells were fixed in 2% PFA prepared in PBS at RT for 10 min. Cells were then directly stained for intracellular lipid droplets or were processed for antigen staining by blocking with PBTG (PBS containing 10% normal goat serum, 1% bovine serum albumin (BSA), 0.1% Triton-X100) at RT for 1-2 h. Cells were then incubated with primary antibodies diluted in PBTG (concentrations above) at 4 °C overnight. Following four washes with PBS-T, cells were stained with AlexaFluor-conjugated secondary antibodies in PBTG at RT for 1 h. Following one wash with PBS-T, cells were incubated with Hoechst nuclear stain diluted in PBS at RT for 15 min, followed by three additional washes with PBS-T and a final wash with PBS. Images were acquired using a 40x oil immersion objective on an inverted Zeiss Axiovert 200 spinning disk confocal microscope using solid-state 491, 561 and 644 nm lasers (Spectral Applied) for excitation for collection with an Andor iXon 512x512 EMCCD camera using MetaMorph software. Acquired images were analyzed as described using Fiji software (Schindelin et al., 2012).

### Western Blot

Cell lysates were prepared in RIPA buffer (50 mM Tris-HCl [pH 8.0], 150 mM NaCl, 0.1% sodium dodecyl sulfate [SDS], 0.5% SDC, 1% NP-40 with addition of cOmplete Mini EDTA-free protease inhibitor tablet [Roche]) and incubated on ice for 30 min with frequent vortexing and then clarified at 14,000 *x g* for 20 min at 4 °C. Protein concentration was determined by bicinchoninic (BCA) protein assay (ThermoFisher Scientific) and samples were resolved on 4-12% Bis-Tris gels (Invitrogen) and 4-20% Tris-Glycine gels (Invitrogen) followed by transfer onto polyvinylidene fluoride (PVDF) membrane (EMD Millipore). Membranes were blocked with 5% milk in PBS-T and incubated with primary antibody at 4 °C overnight in 5% milk PBST. Membranes were washed 3x with PBST and incubated with secondary antibodies. For chemiluminescent readout, the membranes were incubated with HRP-conjugated secondary antibody and exposed using SuperSignal West Pico or Femto PLUS Chemiluminescent Substrate (ThermoFisher Scientific) by film in a dark room.

### Apoptosis assay

Hap1 WT and TMEM41B KO cells were seeded into 24-well plates at 5 x 10^4^ cells/well. The next day, cells were infected with the YFV vaccine strain 17D, UV-inactivated YFV 17D and IAV (ΔNS1) diluted in Opti-MEM for 2 h at 37 °C. Staurosporine (STS) diluted in the culture medium was added to the cells simultaneously. After 2 h cells were washed with Opti-MEM and cultured in fresh medium. After 24 h, detached cells (from supernatant and PBS wash) together with the trypsinized cells were harvested into tubes and samples were prepared on ice for flow cytometry using the Pacific Blue Annexin V Apoptosis Detection Kit with 7-AAD (Biolegend: 640926) according to the manufacturer’s protocol. Cells were subsequently washed twice in binding buffer, followed by fixation in 2% PFA in PBS before subjecting them to flow cytometry using the LSR-II (BD Biosciences). Data was analyzed by FlowJo v10 and graphed by Prism 8 (GraphPad).

### ISG response

HAP1 WT and TMEM41B KO cells were seeded into 24-well plates at 7.5 x 10^4^ cells/well. The next day, cells were infected with the YFV vaccine strain 17D, UV-inactivated YFV 17D and IAV (ΔNS1) diluted in Opti-MEM for 2 h at 37 °C. After 2 h cells were washed with Opti-MEM before adding fresh culture medium. After 24 h at 37 °C, the cells were washed once with PBS, trypsinized, and pelleted before extracting total RNA by using RNeasy Mini Kit (QIAgen: 74104) following the manufacturer’s instructions. Next, 0.2 µg RNA were reverse transcribed into cDNA using random hexamer primers with the Superscript III First-Strand Synthesis System Kit (Invitrogen: 18080051) following the manufacturer’s instructions. Gene expression was quantified by qRT-PCR using PowerUp SYBR Green Master Mix (Applied Biosystems: A25742) and gene-specific primers: RPS11: forward: 5’-GCCGAGACTATCTGCACTAC-3’ and reverse: 5’-ATGTCCAGCCTCAGAACTTC-3’; OAS1: forward: 5’-AGAAATACCCCAGCCAAATCTCT-3’ and reverse: 5’-TGAGGAGCCACCCTTTACCA-3’. The following PCR conditions were used: 56 °C for 2 min and 95 °C for 10 min (initial denaturation); 45 cycles 95 °C for 15 sec, 56 °C for 15 sec and 72 °C for 20 sec (PCR); followed by 95 °C for 10 sec, 65 °C for 10 sec, a slow increase to 95 °C (0.07 °C/sec) and a final cooling to 50 °C (melt curve). The data were analyzed by melt curve analysis for product specificity as well as ΔΔCT analysis for fold changes (after normalization to housekeeping genes) and graphed using Prism 8 (GraphPad).

---

**Figure S1. Related to Figure 1.**
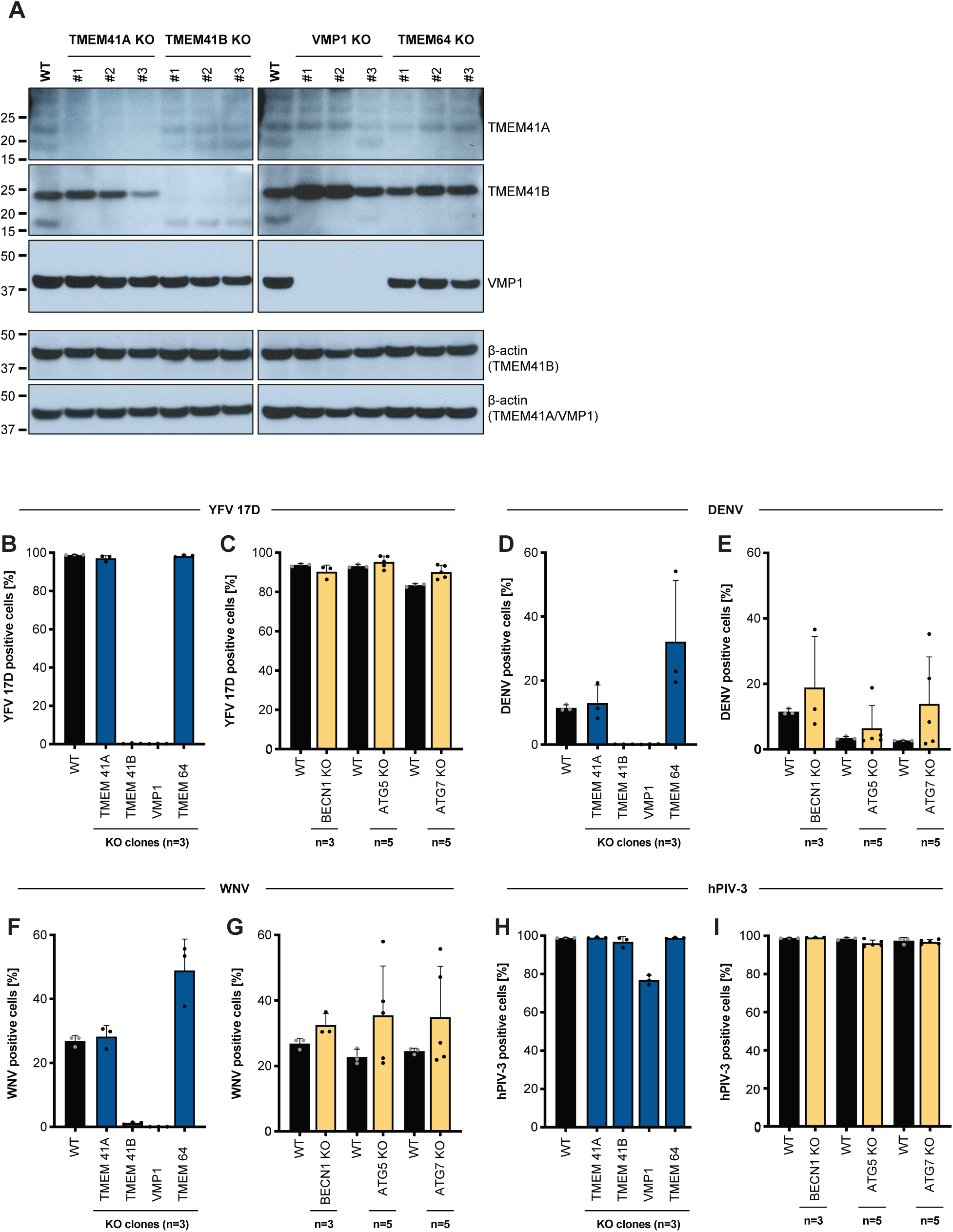
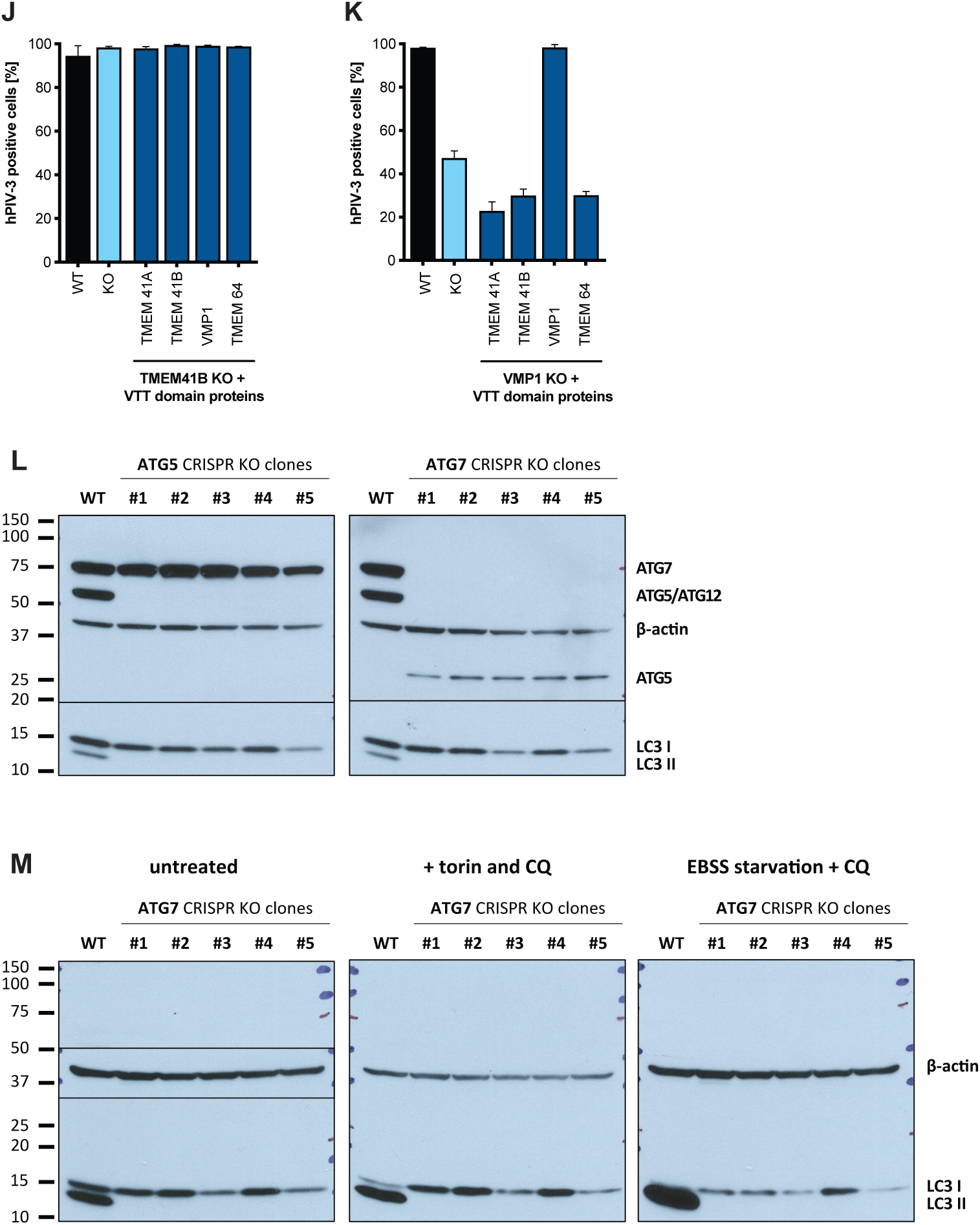
**(A)** Western blots for TMEM41A, TMEM41B, and VMP1 in KO clones lacking individual VTT domain-containing proteins. The expected sizes are indicated by labels on the right. β-actin is shown as loading control. Gene disruption was additionally confirmed by next generation sequencing for each clone. **(B)** HAP1 wildtype (WT) and (n=3) individual knockout (KO) clones for VTT domain-containing proteins infected with YFV 17D (MOI = 0.005 PFU/cell) for 48 h. **(C)** HAP1 WT and (n=3 or 5) individual KO clones for autophagy genes infected with YFV 17D (MOI = 0.005 PFU/cell) for 48 h. **(D-E)** Same as panels B-C but infected with DENV (MOI = 0.1 PFU/cell) for 96 h. **(F-G)** Same as panels B-C but infected with WNV (MOI = 10 PFU/cell) for 72 h. **(H-I)** Same as panels B-C but infected with hPIV-3 (MOI = 0.0.2 IU/cell) for 48 h. **(J)** TMEM41B KO HAP1 cells overexpressing individual VTT domain proteins infected with hPIV-3 (MOI = 0.02 IU/cell) for 48 h. **(K)** VMP1 KO HAP1 cells overexpressing individual VTT domain proteins infected with hPIV-3 (MOI = 0.02 IU/cell) for 72 h. Cells were analyzed by flow cytometry and plotted as a percentage of viral antigen positive cells. Dots in panels B-I represent the average of n=3-5 replicates from individual single cell clones. Error bars in panels J-K are SD of n=3 replicates. **(L)** Western blots for ATG5 and ATG7 in HAP1 WT cells and ATG5 and ATG7 KO clones. The expected sizes are indicated by labels on the right. β-actin is shown as loading control. The anti-ATG5 antibody also recognizes the ATG5/ATG12 conjugate with a higher molecular weight band. Also shown is the differently lipidated LC3 protein (LC3 I/II), indicative of functional autophagy. **(M)** ATG7 KO HAP1 clones untreated (left), treated with Torin 1 (inducer of autophagy) and chloroquine (CQ – block of autophagosome/lysosome fusion and LC3 II turnover) (middle), and serum-starved (induction of autophagy) in Earle’s balanced salt solution (EBSS and treated with CQ) (right) (treatment for 6 h). β-actin is shown as loading control. Functional autophagy and autophagy induction is observed by the appearance of LC3 II, which is absent at baseline in ATG5 and ATG7 KO clones and also upon induction in ATG7 KO clones.

**Figure S2. Related to Figure 2.**
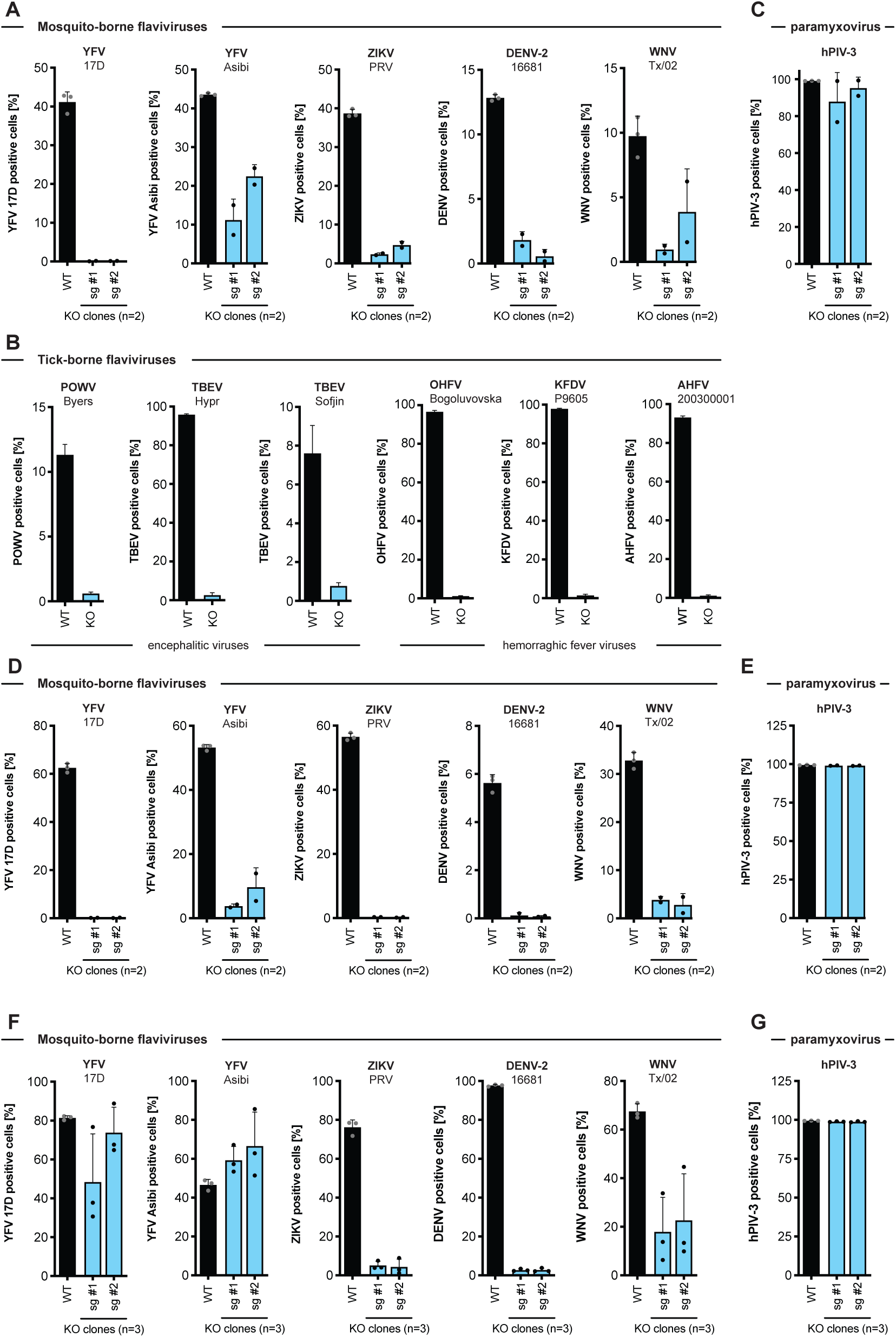
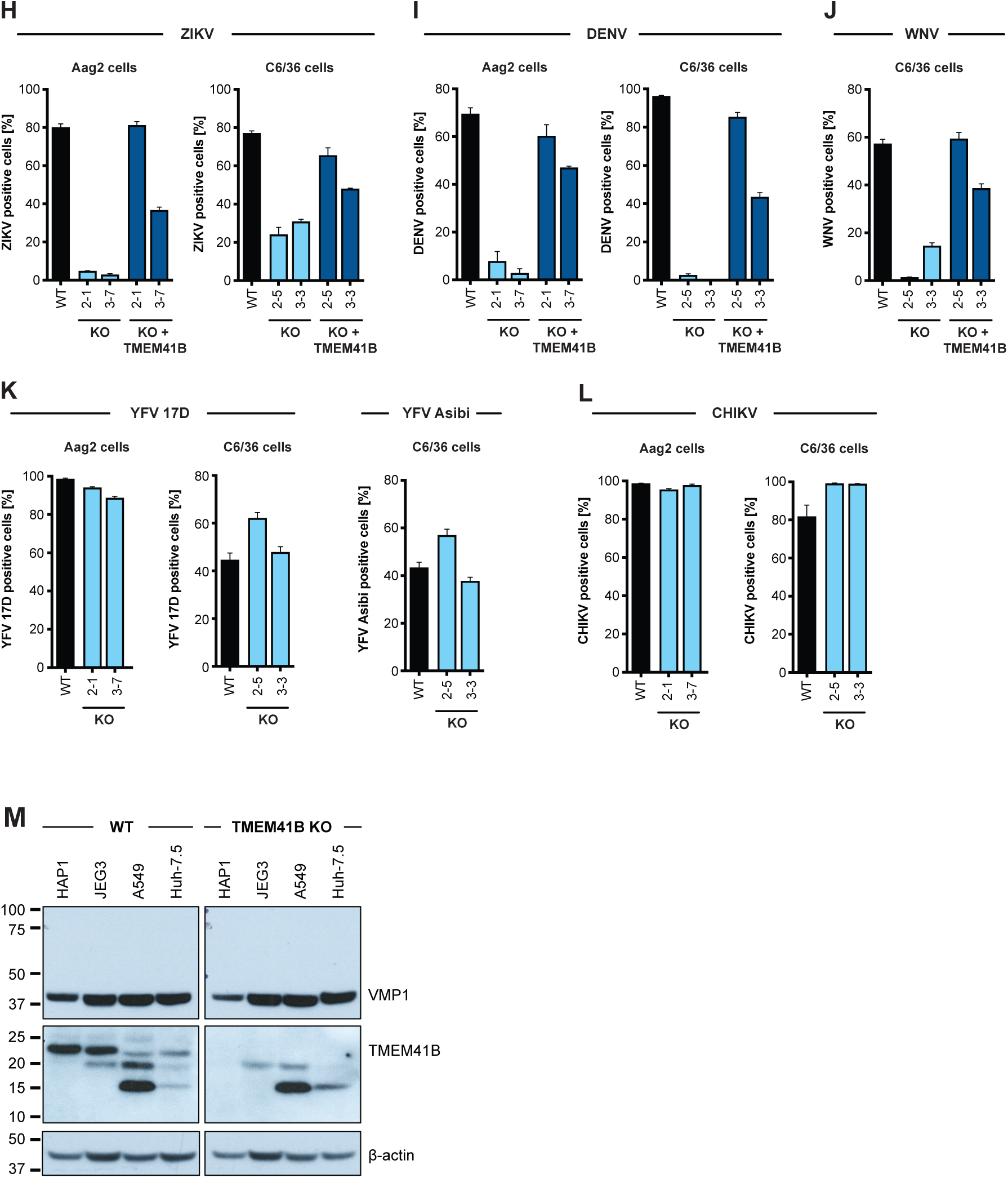
**(A)** WT and TMEM41B KO A549 clones generated with two independent sgRNAs infected with mosquito-borne flaviviruses: YFV 17D, MOI = 0.025 PFU/cell for 48 h; YFV Asibi, MOI = 0.05 PFU/cell for 72 h; ZIKV, MOI = 0.05 PFU/cell for 48 h; DENV-GFP, MOI = 0.1 PFU/cell for 48 h; WNV-GFP, MOI = 1 PFU/cell for 72 h. **(B)** WT and TMEM41B KO A549 clones infected for 48 h with tick-borne flaviviruses at MOIs of 0.02 PFU/cell. **(C)** WT and TMEM41B KO A549 clones infected for 48 h with hPIV-3-GFP, MOI = 0.02 IU/cell. **(D)** WT and TMEM41B KO JEG3 clones generated with two independent sgRNAs infected with mosquito-borne flaviviruses: YFV 17D, MOI = 0.025 PFU/cell for 48 h; YFV Asibi, MOI = 0.05 PFU/cell for 72 h; ZIKV, MOI = 0.025 PFU/cell for 48 h; DENV-GFP, MOI = 0.1 PFU/cell for 72 h; WNV-GFP, MOI = 0.1 PFU/cell for 48 h. **(E)** WT and TMEM41B KO JEG3 clones infected for 48 h with hPIV-3-GFP, MOI = 0.02 IU/cell. **(F)** WT and TMEM41B KO Huh-7.5 clones generated with two independent sgRNAs infected with mosquito-borne flaviviruses: YFV 17D, MOI = 0.0025 PFU/cell for 48 h; YFV Asibi, MOI = 0.0025 PFU/cell for 48 h; ZIKV, MOI = 0.01 PFU/cell for 48 h; DENV-GFP, MOI = 0.01 PFU/cell for 72 h; WNV-GFP, MOI = 0.01 PFU/cell for 48 h. **(G)** WT and TMEM41B KO JEG3 clones infected for 48 h with hPIV-3-GFP, MOI = 0.02 IU/cell. **(H-J)** WT and TMEM41B KO and KO reconstituted Aag2 and C6/36 mosquito cell clones generated with independent sgRNAs infected with: ZIKV, MOI = 0.05 PFU/cell for 72 h; DENV, MOI = 0.2 PFU/cell for 96 h; WNV-GFP, MOI = 10 PFU/cell for 72 h. **(K-L)** Same as H-J but without TMEM41B reconstitution. YFV 17D, MOI = 10 PFU/cell for 72 h; YFV Asibi, MOI = 0.5 PFU/cell for 72 h; CHIKV, MOI = 0.1 PFU/cell for 72 h (Aag2) and 0.05 PFU/cell for 48 h (C6/36). **(M)** Western blot to detect TMEM41B and VMP1 in lysates from indicated cell lines. β-actin is included as a loading control. Panels A, C and D-G, individual dots are the average of n=3 replicates from independent cell clones. Cells were analyzed by flow cytometry and plotted as a percentage of viral antigen positive cells or GFP/RFP positive cells for reporter viruses expressing a fluorescent protein. Individual dot are independent cell clones. Panels A-L, error bars depict SD for three replicates or the indicated number of clones.

**Figure S3. Related to Figure 3.**
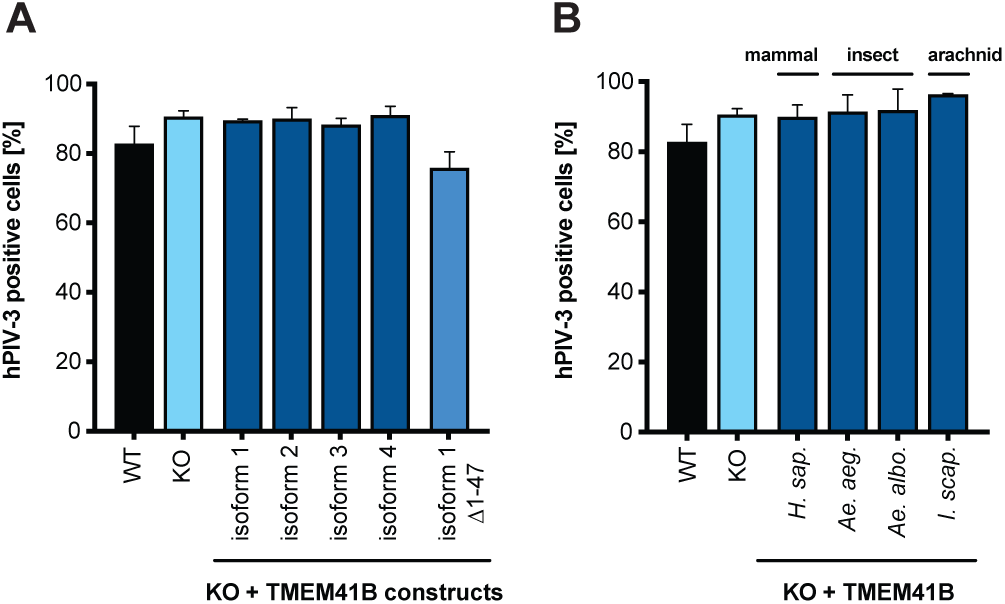
WT, TMEM41B KO, and TMEM41B KO HAP1 cells expressing **(A)** human TMEM41B isoforms or **(B)** mosquito and tick TMEM41B isoforms infected with hPIV-3-GFP MOI = 0.02 IU/cell for 48 h. Cells were analyzed by flow cytometry and plotted as a percentage of GFP positive cells. Error bars depict SD for three replicates.

**Figure S4. Related to Figure 4.**
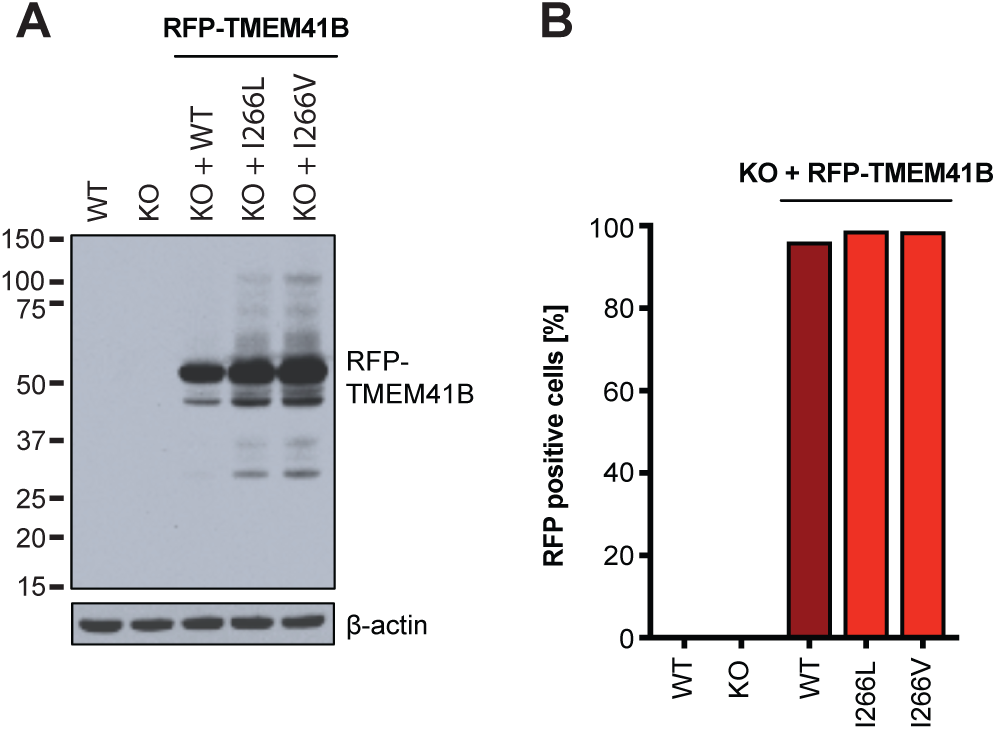
**(A)** Western blot to detect RFP-TMEM41B in HAP1 cell lysates. Predicted size is indicated by the location of the label. β-actin is included as a loading control. (B) Flow cytometry analysis to quantify the percentage of RFP-TMEM41B cells.

**Figure S5. Related to Figure 5.**
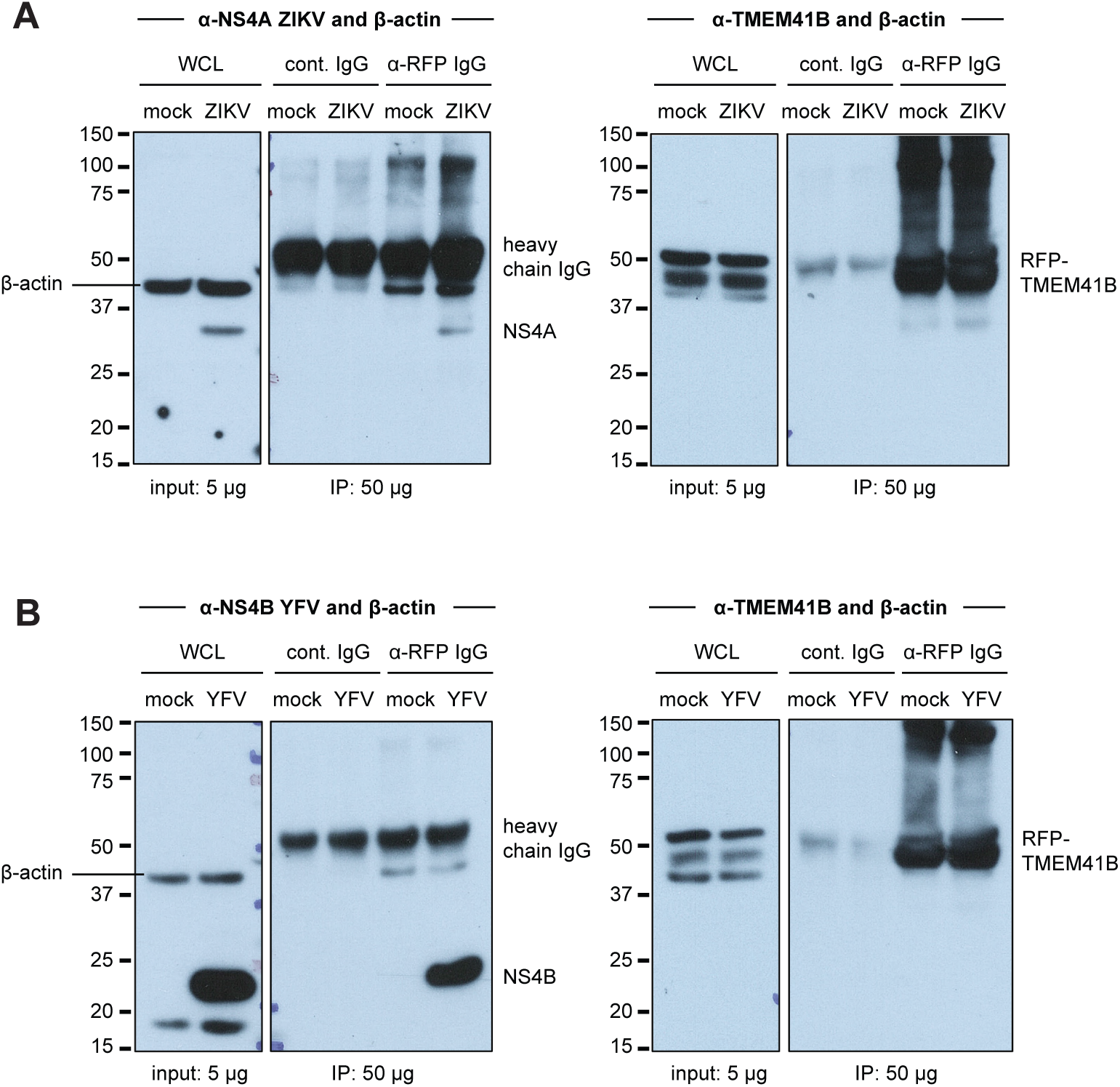
**(A)** Shown are western blots from Figure 5 from ZIKV-infected lysates. Left panel was probed with β-actin to show that similar amounts of whole cell lysate were used as input. Right panel was probed with anti-RFP to show that similar amounts of RFP-TMEM41B were immunoprecipitated from mock and infected lysates but not when using an IgG control antibody. **(B)** Same as (A) with YFV-infected cells. Of note, the β-actin signal detected in the anti-RFP IgG immunopreciptated fraction is likely an unspecific interaction with the RFP or with TMEM41B as it has been shown that the cytoskeleton (e.g., actins) can interact with organelles (e.g., ER).

## REFERENCES

Yellow fever in Africa and the Americas, 2016. Releve epidemiologique hebdomadaire 92, 442–452.

Ahmed, Q.A., and Memish, Z.A. (2017). Yellow fever from Angola and Congo: a storm gathers. Tropical doctor 47, 92–96.

Aktepe, T.E., and Mackenzie, J.M. (2018). Shaping the flavivirus replication complex: It is curvaceous! Cellular microbiology *20*, e12884.

Baggen, J., Persoons, L., Jansen, S., Vanstreels, E., Jacquemyn, M., Jochmans, D., Neyts, J., Dallmeier, K., Maes, P., and Daelemans, D. (2020). Identification of TMEM106B as proviral host factor for SARS-CoV-2. bioRxiv. doi: https://doi.org/10.1101/2020.09.28.316281

Benboudjema, L., Mulvey, M., Gao, Y., Pimplikar, S.W., and Mohr, I. (2003). Association of the herpes simplex virus type 1 Us11 gene product with the cellular kinesin light-chain-related protein PAT1 results in the redistribution of both polypeptides. Journal of virology 77, 9192–9203.

Blight, K.J., McKeating, J.A., and Rice, C.M. (2002). Highly permissive cell lines for subgenomic and genomic hepatitis C virus RNA replication. Journal of virology 76, 13001–13014.

Brady, O.J., and Hay, S.I. (2020). The Global Expansion of Dengue: How Aedes aegypti Mosquitoes Enabled the First Pandemic Arbovirus. Annual review of entomology 65, 191–208.

Brugueras, S., Fernandez-Martinez, B., Martinez-de la Puente, J., Figuerola, J., Porro, T.M., Rius, C., Larrauri, A., and Gomez-Barroso, D. (2020). Environmental drivers, climate change and emergent diseases transmitted by mosquitoes and their vectors in southern Europe: A systematic review. Environmental research 191, 110038.

Cagno, V., Tseligka, E.D., Jones, S.T., and Tapparel, C. (2019). Heparan Sulfate Proteoglycans and Viral Attachment: True Receptors or Adaptation Bias? Viruses 11.

Dalton, K.P., and Rose, J.K. (2001). Vesicular stomatitis virus glycoprotein containing the entire green fluorescent protein on its cytoplasmic domain is incorporated efficiently into virus particles. Virology 279, 414–421.

Dennis, D.T., Nekomoto, T.S., Victor, J.C., Paul, W.S., and Piesman, J. (1998). Reported distribution of Ixodes scapularis and Ixodes pacificus (Acari: Ixodidae) in the United States. Journal of medical entomology 35, 629–638.

Dikic, I., and Elazar, Z. (2018). Mechanism and medical implications of mammalian autophagy. Nature reviews Molecular cell biology 19, 349–364.

Dobler, G. (2010). Zoonotic tick-borne flaviviruses. Veterinary microbiology 140, 221–228.

dos Santos, C.N., Post, P.R., Carvalho, R., Ferreira, II, Rice, C.M., and Galler, R. (1995). Complete nucleotide sequence of yellow fever virus vaccine strains 17DD and 17D-213. Virus research 35, 35–41.

Drappier, M., and Michiels, T. (2015). Inhibition of the OAS/RNase L pathway by viruses. Current opinion in virology 15, 19–26.

Ebel, G.D. (2010). Update on Powassan virus: emergence of a North American tick-borne flavivirus. Annual review of entomology 55, 95–110.

Eisen, R.J., Eisen, L., and Beard, C.B. (2016). County-Scale Distribution of Ixodes scapularis and Ixodes pacificus (Acari: Ixodidae) in the Continental United States. Journal of medical entomology 53, 349–386.

Feuer, R., Mena, I., Pagarigan, R., Slifka, M.K., and Whitton, J.L. (2002). Cell cycle status affects coxsackievirus replication, persistence, and reactivation in vitro. Journal of virology 76, 4430–4440.

Flint, M., McMullan, L.K., Dodd, K.A., Bird, B.H., Khristova, M.L., Nichol, S.T., and Spiropoulou, C.F. (2014). Inhibitors of the tick-borne, hemorrhagic fever-associated flaviviruses. Antimicrobial agents and chemotherapy 58, 3206–3216.

Garcia-Sastre, A., Egorov, A., Matassov, D., Brandt, S., Levy, D.E., Durbin, J.E., Palese, P., and Muster, T. (1998). Influenza A virus lacking the NS1 gene replicates in interferon-deficient systems. Virology 252, 324–330.

Gusho, E., Baskar, D., and Banerjee, S. (2020). New advances in our understanding of the "unique" RNase L in host pathogen interaction and immune signaling. Cytokine 133, 153847.

Haeussler, M., Schonig, K., Eckert, H., Eschstruth, A., Mianne, J., Renaud, J.B., Schneider-Maunoury, S., Shkumatava, A., Teboul, L., Kent, J., et al. (2016). Evaluation of off-target and on-target scoring algorithms and integration into the guide RNA selection tool CRISPOR. Genome biology 17, 148.

Hills, S.L., Fischer, M., and Petersen, L.R. (2017). Epidemiology of Zika Virus Infection. The Journal of infectious diseases 216, S868–S874.

Hoffmann, H.H., Schneider, W.M., Blomen, V.A., Scull, M.A., Hovnanian, A., Brummelkamp, T.R., and Rice, C.M. (2017). Diverse Viruses Require the Calcium Transporter SPCA1 for Maturation and Spread. Cell host & microbe 22, 460–470 e465.

Ishii, K.J., and Akira, S. (2005). Innate immune recognition of nucleic acids: beyond toll-like receptors. International journal of cancer 117, 517–523.

Jae, L.T., Raaben, M., Herbert, A.S., Kuehne, A.I., Wirchnianski, A.S., Soh, T.K., Stubbs, S.H., Janssen, H., Damme, M., Saftig, P., et al. (2014). Virus entry. Lassa virus entry requires a trigger-induced receptor switch. Science 344, 1506–1510.

Kang, R., Zeh, H.J., Lotze, M.T., and Tang, D. (2011). The Beclin 1 network regulates autophagy and apoptosis. Cell death and differentiation 18, 571–580.

Kang, Z.B., Moutsatsos, I., Moretti, F., Bergman, P., Zhang, X., Nyfeler, B., and Antczak, C. (2020). Fluopack screening platform for unbiased cellular phenotype profiling. Scientific reports 10, 2097.

Ke, P.Y. (2018). The Multifaceted Roles of Autophagy in Flavivirus-Host Interactions. International journal of molecular sciences 19.

Kelley, L.A., Mezulis, S., Yates, C.M., Wass, M.N., and Sternberg, M.J. (2015). The Phyre2 web portal for protein modeling, prediction and analysis. Nature protocols 10, 845–858.

Kramer, L.D., Ciota, A.T., and Kilpatrick, A.M. (2019). Introduction, Spread, and Establishment of West Nile Virus in the Americas. Journal of medical entomology 56, 1448–1455.

Lee, C.Y., and Ng, L.F.P. (2018). Zika virus: from an obscurity to a priority. Microbes and infection 20, 635–645.

Li, W., Xu, H., Xiao, T., Cong, L., Love, M.I., Zhang, F., Irizarry, R.A., Liu, J.S., Brown, M., and Liu, X.S. (2014). MAGeCK enables robust identification of essential genes from genome-scale CRISPR/Cas9 knockout screens. Genome biology 15, 554.

Liu, D., Ji, J., Ndongwe, T.P., Michailidis, E., Rice, C.M., Ralston, R., and Sarafianos, S.G. (2015). Fast hepatitis C virus RNA elimination and NS5A redistribution by NS5A inhibitors studied by a multiplex assay approach. Antimicrobial agents and chemotherapy 59, 3482–3492.

Marceau, C.D., Puschnik, A.S., Majzoub, K., Ooi, Y.S., Brewer, S.M., Fuchs, G., Swaminathan, K., Mata, M.A., Elias, J.E., Sarnow, P., et al. (2016). Genetic dissection of Flaviviridae host factors through genome-scale CRISPR screens. Nature 535, 159–163.

McGee, C.E., Shustov, A.V., Tsetsarkin, K., Frolov, I.V., Mason, P.W., Vanlandingham, D.L., and Higgs, S. (2010). Infection, dissemination, and transmission of a West Nile virus green fluorescent protein infectious clone by Culex pipiens quinquefasciatus mosquitoes. Vector borne and zoonotic diseases 10, 267–274.

McPherson, M., Garcia-Garcia, A., Cuesta-Valero, F.J., Beltrami, H., Hansen-Ketchum, P., MacDougall, D., and Ogden, N.H. (2017). Expansion of the Lyme Disease Vector Ixodes Scapularis in Canada Inferred from CMIP5 Climate Projections. Environmental health perspectives 125, 057008.

Medlock, J.M., Hansford, K.M., Bormane, A., Derdakova, M., Estrada-Pena, A., George, J.C., Golovljova, I., Jaenson, T.G., Jensen, J.K., Jensen, P.M., et al. (2013). Driving forces for changes in geographical distribution of Ixodes ricinus ticks in Europe. Parasites & vectors 6, 1.

Mendez, E., Ruggli, N., Collett, M.S., and Rice, C.M. (1998). Infectious bovine viral diarrhea virus (strain NADL) RNA from stable cDNA clones: a cellular insert determines NS3 production and viral cytopathogenicity. Journal of virology 72, 4737–4745.

Mizushima, N., Noda, T., Yoshimori, T., Tanaka, Y., Ishii, T., George, M.D., Klionsky, D.J., Ohsumi, M., and Ohsumi, Y. (1998). A protein conjugation system essential for autophagy. Nature 395, 395–398.

Moretti, F., Bergman, P., Dodgson, S., Marcellin, D., Claerr, I., Goodwin, J.M., DeJesus, R., Kang, Z., Antczak, C., Begue, D., et al. (2018). TMEM41B is a novel regulator of autophagy and lipid mobilization. EMBO reports 19.

Morishita, H., Zhao, Y.G., Tamura, N., Nishimura, T., Kanda, Y., Sakamaki, Y., Okazaki, M., Li, D., and Mizushima, N. (2019). A critical role of VMP1 in lipoprotein secretion. eLife 8.

Morita, K., Hama, Y., and Mizushima, N. (2019). TMEM41B functions with VMP1 in autophagosome formation. Autophagy 15, 922–923.

Muller, P., Engeler, L., Vavassori, L., Suter, T., Guidi, V., Gschwind, M., Tonolla, M., and Flacio, E. (2020). Surveillance of invasive Aedes mosquitoes along Swiss traffic axes reveals different dispersal modes for Aedes albopictus and Ae. japonicus. PLoS neglected tropical diseases 14, e0008705.

Neufeldt, C.J., Cortese, M., Acosta, E.G., and Bartenschlager, R. (2018). Rewiring cellular networks by members of the Flaviviridae family. Nature reviews Microbiology 16, 125–142.

Paul, D., and Bartenschlager, R. (2015). Flaviviridae Replication Organelles: Oh, What a Tangled Web We Weave. Annual review of virology 2, 289–310.

Popovic, D., Akutsu, M., Novak, I., Harper, J.W., Behrends, C., and Dikic, I. (2012). Rab GTPase-activating proteins in autophagy: regulation of endocytic and autophagy pathways by direct binding to human ATG8 modifiers. Molecular and cellular biology 32, 1733–1744.

Popovic, D., and Dikic, I. (2014). TBC1D5 and the AP2 complex regulate ATG9 trafficking and initiation of autophagy. EMBO reports 15, 392–401.

Rajah, M.M., Monel, B., and Schwartz, O. (2020). The entanglement between flaviviruses and ER-shaping proteins. PLoS pathogens 16, e1008389.

Realegeno, S., Puschnik, A.S., Kumar, A., Goldsmith, C., Burgado, J., Sambhara, S., Olson, V.A., Carroll, D., Damon, I., Hirata, T., et al. (2017). Monkeypox Virus Host Factor Screen Using Haploid Cells Identifies Essential Role of GARP Complex in Extracellular Virus Formation. Journal of virology 91.

Riblett, A.M., Blomen, V.A., Jae, L.T., Altamura, L.A., Doms, R.W., Brummelkamp, T.R., and Wojcechowskyj, J.A. (2016). A Haploid Genetic Screen Identifies Heparan Sulfate Proteoglycans Supporting Rift Valley Fever Virus Infection. Journal of virology 90, 1414-1423.

Roehrig, J.T., Layton, M., Smith, P., Campbell, G.L., Nasci, R., and Lanciotti, R.S. (2002). The emergence of West Nile virus in North America: ecology, epidemiology, and surveillance. Current topics in microbiology and immunology 267, 223–240.

Ropolo, A., Grasso, D., Pardo, R., Sacchetti, M.L., Archange, C., Lo Re, A., Seux, M., Nowak, J., Gonzalez, C.D., Iovanna, J.L., et al. (2007). The pancreatitis-induced vacuole membrane protein 1 triggers autophagy in mammalian cells. The Journal of biological chemistry 282, 37124–37133.

Savidis, G., McDougall, W.M., Meraner, P., Perreira, J.M., Portmann, J.M., Trincucci, G., John, S.P., Aker, A.M., Renzette, N., Robbins, D.R., et al. (2016). Identification of Zika Virus and Dengue Virus Dependency Factors using Functional Genomics. Cell reports 16, 232–246.

Schindelin, J., Arganda-Carreras, I., Frise, E., Kaynig, V., Longair, M., Pietzsch, T., Preibisch, S., Rueden, C., Saalfeld, S., Schmid, B., et al. (2012). Fiji: an open-source platform for biological-image analysis. Nature methods 9, 676–682.

Schneider, W.M., Luna, J.M., Hoffmann, H.H., Sánchez-Rivera, F.J., Leal, A.A., Ashbrook, A.W., Le Pen, J., Michailidis, E., Ricardo-Lax, I., Peace, A., et al. (2020). Genome-scale identification of SARS-CoV-2 and pan-coronavirus host factor networks. bioRxiv. doi: https://doi.org/10.1101/2020.10.07.326462

Schoggins, J.W., Dorner, M., Feulner, M., Imanaka, N., Murphy, M.Y., Ploss, A., and Rice, C.M. (2012). Dengue reporter viruses reveal viral dynamics in interferon receptor-deficient mice and sensitivity to interferon effectors in vitro. Proceedings of the National Academy of Sciences of the United States of America 109, 14610–14615.

Schoggins, J.W., MacDuff, D.A., Imanaka, N., Gainey, M.D., Shrestha, B., Eitson, J.L., Mar, K.B., Richardson, R.B., Ratushny, A.V., Litvak, V., et al. (2014). Pan-viral specificity of IFN-induced genes reveals new roles for cGAS in innate immunity. Nature 505, 691–695.

Schwartz, S.L., and Conn, G.L. (2019). RNA regulation of the antiviral protein 2’-5’-oligoadenylate synthetase. Wiley interdisciplinary reviews RNA 10, e1534.

Shalem, O., Sanjana, N.E., Hartenian, E., Shi, X., Scott, D.A., Mikkelson, T., Heckl, D., Ebert, B.L., Root, D.E., Doench, J.G., et al. (2014). Genome-scale CRISPR-Cas9 knockout screening in human cells. Science 343, 84–87.

Shan, C., Xie, X., Muruato, A.E., Rossi, S.L., Roundy, C.M., Azar, S.R., Yang, Y., Tesh, R.B., Bourne, N., Barrett, A.D., et al. (2016). An Infectious cDNA Clone of Zika Virus to Study Viral Virulence, Mosquito Transmission, and Antiviral Inhibitors. Cell host & microbe 19, 891–900.

Shi, X., van Mierlo, J.T., French, A., and Elliott, R.M. (2010). Visualizing the replication cycle of bunyamwera orthobunyavirus expressing fluorescent protein-tagged Gc glycoprotein. Journal of virology 84, 8460–8469.

Shoemaker, C.J., Huang, T.Q., Weir, N.R., Polyakov, N.J., Schultz, S.W., and Denic, V. (2019). CRISPR screening using an expanded toolkit of autophagy reporters identifies TMEM41B as a novel autophagy factor. PLoS biology 17, e2007044.

Sidjanin, D.J., Park, A.K., Ronchetti, A., Martins, J., and Jackson, W.T. (2016). TBC1D20 mediates autophagy as a key regulator of autophagosome maturation. Autophagy 12, 1759–1775.

Sklan, E.H., Serrano, R.L., Einav, S., Pfeffer, S.R., Lambright, D.G., and Glenn, J.S. (2007). TBC1D20 is a Rab1 GTPase-activating protein that mediates hepatitis C virus replication. The Journal of biological chemistry 282, 36354–36361.

Tabara, L.C., and Escalante, R. (2016). VMP1 Establishes ER-Microdomains that Regulate Membrane Contact Sites and Autophagy. PloS one 11, e0166499.

Tanida, I., Mizushima, N., Kiyooka, M., Ohsumi, M., Ueno, T., Ohsumi, Y., and Kominami, E. (1999). Apg7p/Cvt2p: A novel protein-activating enzyme essential for autophagy. Molecular biology of the cell 10, 1367–1379.

Tanida, I., Ueno, T., and Kominami, E. (2008). LC3 and Autophagy. Methods in molecular biology 445, 77–88.

Tanida, I., and Waguri, S. (2010). Measurement of autophagy in cells and tissues. Methods in molecular biology 648, 193–214.

Tsetsarkin, K., Higgs, S., McGee, C.E., De Lamballerie, X., Charrel, R.N., and Vanlandingham, D.L. (2006). Infectious clones of Chikungunya virus (La Reunion isolate) for vector competence studies. Vector borne and zoonotic diseases 6, 325–337.

Wang, R., Simoneau, C.R., Kulsuptrakul, J., Bouhaddou, M., Travisano, K., Hayashi, J.M., Carlson-Stevermer, J., Oki, J., Holden, K., Krogan, N.J., et al. (2020). Functional genomic screens identify human host factors for SARS-CoV-2 and common cold coronaviruses. bioRxiv. doi: https://doi.org/10.1101/2020.09.24.312298

Wang, X., Li, M., Zheng, H., Muster, T., Palese, P., Beg, A.A., and Garcia-Sastre, A. (2000). Influenza A virus NS1 protein prevents activation of NF-kappaB and induction of alpha/beta interferon. Journal of virology 74, 11566–11573.

Yau, E.H., and Rana, T.M. (2018). Next-Generation Sequencing of Genome-Wide CRISPR Screens. Methods in molecular biology 1712, 203–216.

Zaki, A.M. (1997). Isolation of a flavivirus related to the tick-borne encephalitis complex from human cases in Saudi Arabia. Transactions of the Royal Society of Tropical Medicine and Hygiene 91, 179–181.

Zhang, L., Bukreyev, A., Thompson, C.I., Watson, B., Peeples, M.E., Collins, P.L., and Pickles, R.J. (2005). Infection of ciliated cells by human parainfluenza virus type 3 in an in vitro model of human airway epithelium. Journal of virology 79, 1113–1124.

Zhang, R., Miner, J.J., Gorman, M.J., Rausch, K., Ramage, H., White, J.P., Zuiani, A., Zhang, P., Fernandez, E., Zhang, Q., et al. (2016). A CRISPR screen defines a signal peptide processing pathway required by flaviviruses. Nature 535, 164–168.

Zhao, Y.G., Chen, Y., Miao, G., Zhao, H., Qu, W., Li, D., Wang, Z., Liu, N., Li, L., Chen, S., et al. (2017). The ER-Localized Transmembrane Protein EPG-3/VMP1 Regulates SERCA Activity to Control ER-Isolation Membrane Contacts for Autophagosome Formation. Molecular cell 67, 974–989 e976.

